# Stem cells tightly regulate dead cell clearance to maintain tissue fitness

**DOI:** 10.1101/2023.05.22.541773

**Authors:** Katherine S Stewart, Kevin AU Gonzales, Shaopeng Yuan, Matthew T Tierney, Alain R Bonny, Yihao Yang, Nicole R Infarinato, Christopher J Cowley, John M Levorse, Hilda Amalia Pasolli, Sourav Ghosh, Carla V Rothlin, Elaine Fuchs

## Abstract

Macrophages and dendritic cells have long been appreciated for their ability to migrate to and engulf dying cells and debris, including some of the billions of cells that are naturally eliminated from our body daily. However, a substantial number of these dying cells are cleared by ‘non-professional phagocytes’, local epithelial cells that are critical to organismal fitness. How non-professional phagocytes sense and digest nearby apoptotic corpses while still performing their normal tissue functions is unclear. Here, we explore the molecular mechanisms underlying their multifunctionality. Exploiting the cyclical bouts of tissue regeneration and degeneration during the hair cycle, we show that stem cells can transiently become non-professional phagocytes when confronted with dying cells. Adoption of this phagocytic state requires both local lipids produced by apoptotic corpses to activate RXRα, and tissue-specific retinoids for RARγ activation. This dual factor dependency enables tight regulation of the genes requisite to activate phagocytic apoptotic clearance. The tunable phagocytic program we describe here offers an effective mechanism to offset phagocytic duties against the primary stem cell function of replenishing differentiated cells to preserve tissue integrity during homeostasis. Our findings have broad implications for other non-motile stem or progenitor cells which experience cell death in an immune-privileged niche.

## Main

Adult homeostatic tissue maintenance necessitates the turnover and replacement of billions of cells daily ^1,2^. Tissue rejuvenation is achieved by stem cells, which balance proliferation and differentiation through carefully charted programs. However, when cells within a tissue die, they must be removed quickly to maintain immune and tissue homeostasis. Clearance of apoptotic cells (‘efferocytosis’) is the provenance of both professional (immune cells) and non-professional (epithelial and mesenchymal cells) phagocytes^2-5^. In developmental, homeostatic, and pathological contexts, phagocytic cells detect and engulf apoptotic corpses. Failure to do so results in secondary necrosis, which can result in many inflammatory and/or degenerative pathologies^1,6-8^.

Dead cell clearance is a step-wise process initiated when apoptotic corpses transmit “find-me” signals, which are then recognized by clearance receptors on phagocytic cells^2-6^. The sensing mechanisms have been well-described for motile professional phagocytes, such as macrophages and dendritic cells, which are defined by their constitutive phagocytic functions, but remain less clear for non-motile non-professional phagocytes^4^. Nonetheless, engagement of clearance receptors in most if not all phagocytes converges on downstream activation of the Elmo-Dock-Rac pathway to mediate actin cytoskeleton rearrangement and facilitate uptake of apoptotic bodies^2-6^. Subsequent digestion of apoptotic corpses occurs via phagosome maturation and fusion with lysosomes, wherein corpse-derived materials are degraded^2-6,9^.

How non-motile, non-professional phagocytes regulate apoptotic cell clearance in tissues remains unclear. The issue is an important one as in contrast to their immune counterparts, non-professional phagocytes have other dedicated roles in tissue function. We hypothesize that they are able to transiently and expeditiously activate phagocytosis and then dial it down again to return to their normal tissue tasks. Here, we explore this in the context of a dynamic tissue setting by focusing on the synchronized hair cycles of regenerative growth (anagen), degeneration (catagen), and quiescence (telogen) (**Fig. 1a**).

**Fig. 1.**
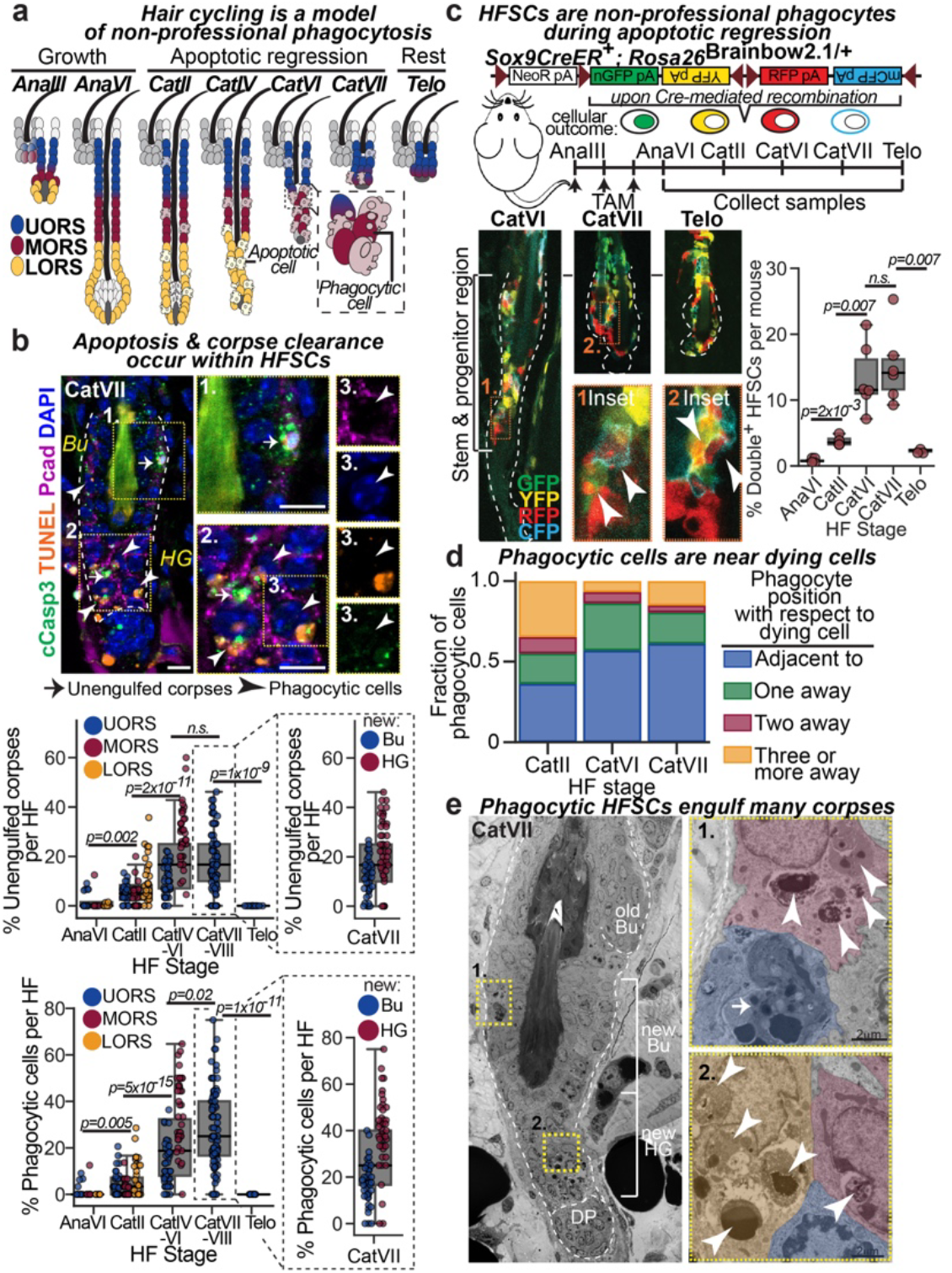
Apoptotic corpses are cleared by neighbouring hair follicle stem cells in late catagen. **a,** Schematic of mammalian hair cycle depicting the old HFSC niche (bulge) in grey and active niche in blue. Ana, anagen; Cat, catagen; Telo, telogen; UORS, upper ORS (cells 1-15 containing HFSCs); MORS, middle ORS (cells 16-30); LORS, lower ORS (cells 31+). **b,** Top, Sagittal skin sections showing apoptic cells (cCasp3^+^TUNEL^+^) encased by upper ORS catagen cells (cell membrane delineated by P-cadherin). DAPI, DNA. Insets show dying cells (arrows), adjacent to phagocytic cells (arrowheads). Quantifications shown below. 10-20 follicles per mouse, *n*=4-6 mice per stage. Bu, bulge (HFSCs); HG, hair germ (early progenitor cells). Dashed lines denote dermo-epithelial border. Scale bar 10um. **c,** Top, strategy to identify functional phagocytic HF stem and progenitor cells. Bottom left, immunofluorescence images of single^+^ (containing no corpses) and double^+^ (phagocytic) HF stem and progenitor cells in the upper ORS. Insets show higher magnification of double^+^ cells that arise from engulfing apoptotic corpses of a different colour (white arrowheads). Bottom right, FACS quantifications. *n*=4-6 mice per stage. **d,** Quantifications across the hair cycle of phagocytic cell position with respect to nearest apoptotic cell. 10-20 follicles per mouse, *n*=4-6 mice per stage. **e,** Ultrastructural images showing multiple apoptotic corpses (white arrowheads) inside late catagen HF stem and progenitor populations. Boxed regions are magnified at right. Pseudocoloured individual cells highlight the presence of phagocytic stem (1.) and progenitor (2.) cells containing engulfed corpses (white arrowheads) near apoptosing cells (white arrow). DP, dermal papilla. All quantifications, multiple pairwise Student’s *T*-Tests, p-values indicated. n.s. not significant (p>0.05).

The hair follicle stem cells (HFSCs) that fuel this cycle reside in a niche (‘bulge’) within the HF. During the early growth phase, they proliferate to generate the outer root sheath (ORS) and the differentiating hair lineages encased therein. The upper ORS becomes quiescent early, setting aside stem and progenitor cells to form a new bulge for the next hair cycle, while the lower ORS contain short-lived progeny that contribute to hair growth.^10,11,12^ During catagen, terminal differentiation of the hair lineages and apoptosis within the ORS^13^ converge with contractile forces from the surrounding dermal sheath^14^ to drive HF involution. Notably, during the regressive phase, apoptotic cells in the lower ORS are cleared by neighbouring epithelial cells^13,15^, but how this occurs remains unclear.

### Hair follicle stem and progenitor cells engulf corpses during physiological apoptosis

To elucidate the spatiotemporal pattern of apoptotic cell clearance across the hair cycle, we used immunofluorescence microscopy to identify dying cells, which are positive for cleaved caspase-3 (Casp3^+^) and DNA damage (terminal-uridine-nick-end-labelling, TUNEL^+^), and monitored their engulfment (TUNEL^+^ apoptotic body inside a healthy P-cadherin^+^ ORS cell with a normal nucleus) (**Fig. 1b**). Dying HF cell numbers increased progressively throughout the catagen ORS. As previously reported, apoptotic cell death was high in the lower ORS.^6,9,12^ That said, apoptotic cells were also found within the upper ORS, previously assumed to be unaffected^10,13^.

In agreement with a prior report^13^, CD45+ professional phagocytes showed no signs of phagocytic activity at this time (**Extended data Fig. 1a**). By contrast, healthy basal ORS epithelial progenitors displayed numerous TUNEL^+^ condensed apoptotic bodies within their cytoplasm, indicative of phagocytosis (**Fig. 1b**). The timing and patterning of phagocytic cells paralleled that of dying cells. Phagocytic HF cells were in equal proportion to apoptotic cells early in catagen (CatII) but increased to roughly twice as many by late catagen (**Extended data Fig. 1b**). Notably, at the end of catagen, engulfed apoptotic corpses were found entirely within the HF stem and progenitor populations responsible for driving the next hair cycle.

To directly test the ability of HFSCs to clear apoptotic corpses, we devised a strategy to detect bona-fide phagocytic cells within catagen-phase HFs. By activating Cre-recombinase in mid-growth HFs of *Sox9CreER^+^*; *R26-Brainbow2.1^fl/+^* mice, we specifically labeled the upper ORS compartment with one of four fluorophores (nuclear GFP, cytoplasmic YFP or RFP, and membrane CFP). Tracing these stochastically labelled stem and progenitor cells across the hair cycle, we visualized apoptotic corpse engulfment of one colour by phagocytic cells of another colour by both immunofluorescence imaging and fluorescent-activated cell sorting (**Fig. 1c**). Wherever cell death was negligible, i.e. at the end of anagen and again during telogen, double-positive cells were largely absent; however, double-positive cells rose markedly during the destructive phase, peaking in late catagen.

Clearance of apoptotic corpses within the mature tissue appeared to necessitate a close spatial relation between the nonmotile phagocyte and the dying cell, as evidenced by nearest apoptotic neighbour analyses of phagocytic cells (**Fig. 1b,d, Extended data Fig. 1c**). Indeed, a majority of phagocytic ORS cells were found either directly adjacent to or one cell body away from an apoptotic cell. Consistent with rapid apoptotic cell recognition and clearance, few stand-alone apoptotic cells were identified per follicle, with many more engulfed apoptotic bodies detected inside healthy basal ORS epithelial progenitors (**Fig. 1b,e**). Indeed, TEM images of HFs in late catagen (CatVII) revealed an average of 2-3 apoptotic corpses per stem or progenitor cell (**Fig. 1d, Extended data Fig. 1d**), suggestive of multiple rounds of engulfment. These findings exposed HFSCs as a cohort of non-professional phagocytes, with important roles in both tissue destruction and regeneration.

### Hair follicle stem cells adopt a transient phagocytic program characterized by *Mertk* expression

The finding that HFSCs are important contributors to apoptotic cell clearance during catagen prompted us to wonder how their phagocytic activity is controlled to prevent elimination of the stem cell pool. To answer this question, we first turned to single-cell transcriptomic profiling of HFSCs across the hair cycle to see if the program might be regulated transcriptionally. Applying the Leiden algorithm, we identified 7 cell clusters which corresponded to hair cycle stage and anatomic location (**Fig. 2a**, **Extended data Fig. 2**). To determine the transcriptional shifts that accompany the transition from HF growth to regression, we performed gene set enrichment analysis on differentially expressed genes from the end of the growth phase (AnaVI) to mid-late destructive phase (CatVI-VII). Notably, the catagen-phase stem and progenitor cells displayed a hair cycle-dependent significant enrichment of gene ontology terms related to a phagocytic state, including apoptotic cell clearance, actin cytoskeletal rearrangement, phagocytic cup formation and trafficking to the lysosome (**Fig. 2b, Extended data Fig. 3a**).

**Fig. 2.**
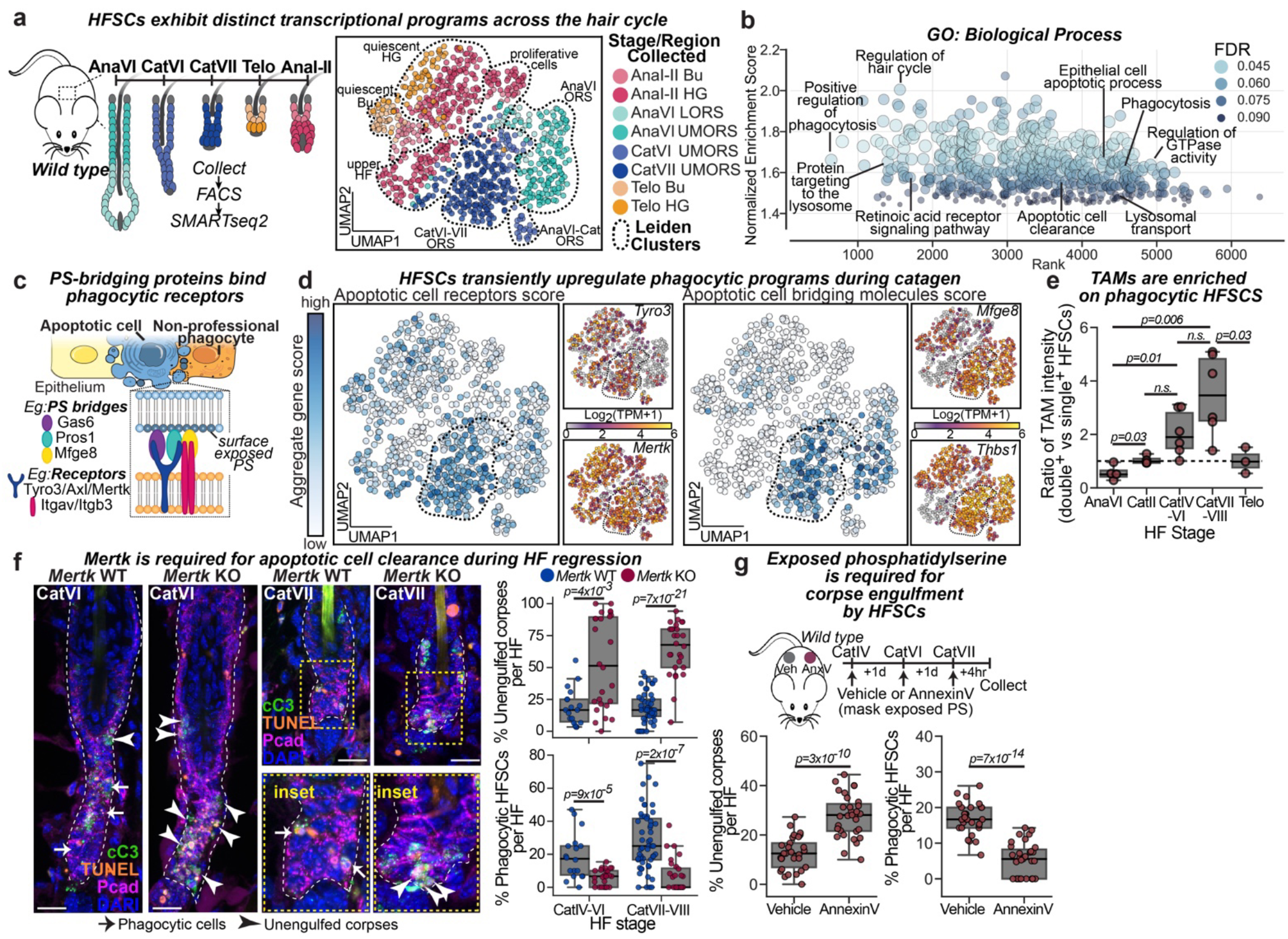
Transient upregulation of phagocytic machinery mediates apoptotic cell clearance in catagen. **a,** Left, strategy to profile hair follicle (HF) epithelial cells across the hair cycle using single-cell RNA-sequencing. Abbreviations as in Fig. 1. Right, UMAP representation and Leiden clustering of single-cell transcriptomes from pooled FACS-isolated CD34^low^ integrin^+^ lower ORS and CD34^high^ integrin^+^ upper ORS cells from wild-type mice. Cells are colour-coded by hair cycle stage and region, and Leiden clusters are encased by black dashed lines. **b,** Gene set enrichment analysis for genes upregulated in catagen versus late anagen; GO, gene ontology. **c,** Schematic depicting apoptotic cell recognition and tethering by neighbouring phagocyte. **d,** UMAP representation of single-cell transcriptomes coloured by aggregate gene set score, with insets showing the relative expression levels (log2[TPM+1]) of example genes from each gene set. The catagen cluster is outlined. Left, Apoptotic cell receptor genes. Right, Apoptotic cell ligands. **e,** *Tyro3*/*Axl*/*Mertk* (TAM)-family phagocytic receptor expression in FACS-purified *Sox9CreER^+^ Brainbow2.1* fluorophore HFSCs that are double^+^ (containing an apoptotic corpse) versus single^+^ (no corpses) per mouse across the hair cycle, *n*=4-6 mice per stage. **f,** Left, Sagittal sections of catagen HFs from wild type (WT) versus constitutive *Mertk* knockout (KO) mice; boxed regions are magnified at right; cC3, cleaved caspase 3; Pcad, p-cadherin; scale bar 20um. Right, Quantifications of unengulfed apoptotic corpses (Top) and corpse-containing phagocytic HFSCs (Bottom). 20 follicles (CatIV-VI) and 65 follicles (CatVII-VII) quantified, *n*=3 mice per stage. **g,** Top, strategy to block corpse-exposed phosphatidylserine (PS) via intradermal injection of soluble AnnexinV (AnxV) during catagen. Contralateral vehicle injections as internal control. Quantifications of unengulfed apoptotic corpses (Bottom left) and corpse-containing phagocytic HFSCs (Bottom right). 5 follicles quantified per mouse, *n*=4 mice. All quantifications, multiple pairwise Student’s *T*-Tests, p-values indicated. n.s. not significant (p>0.05).

**Fig. 3.**
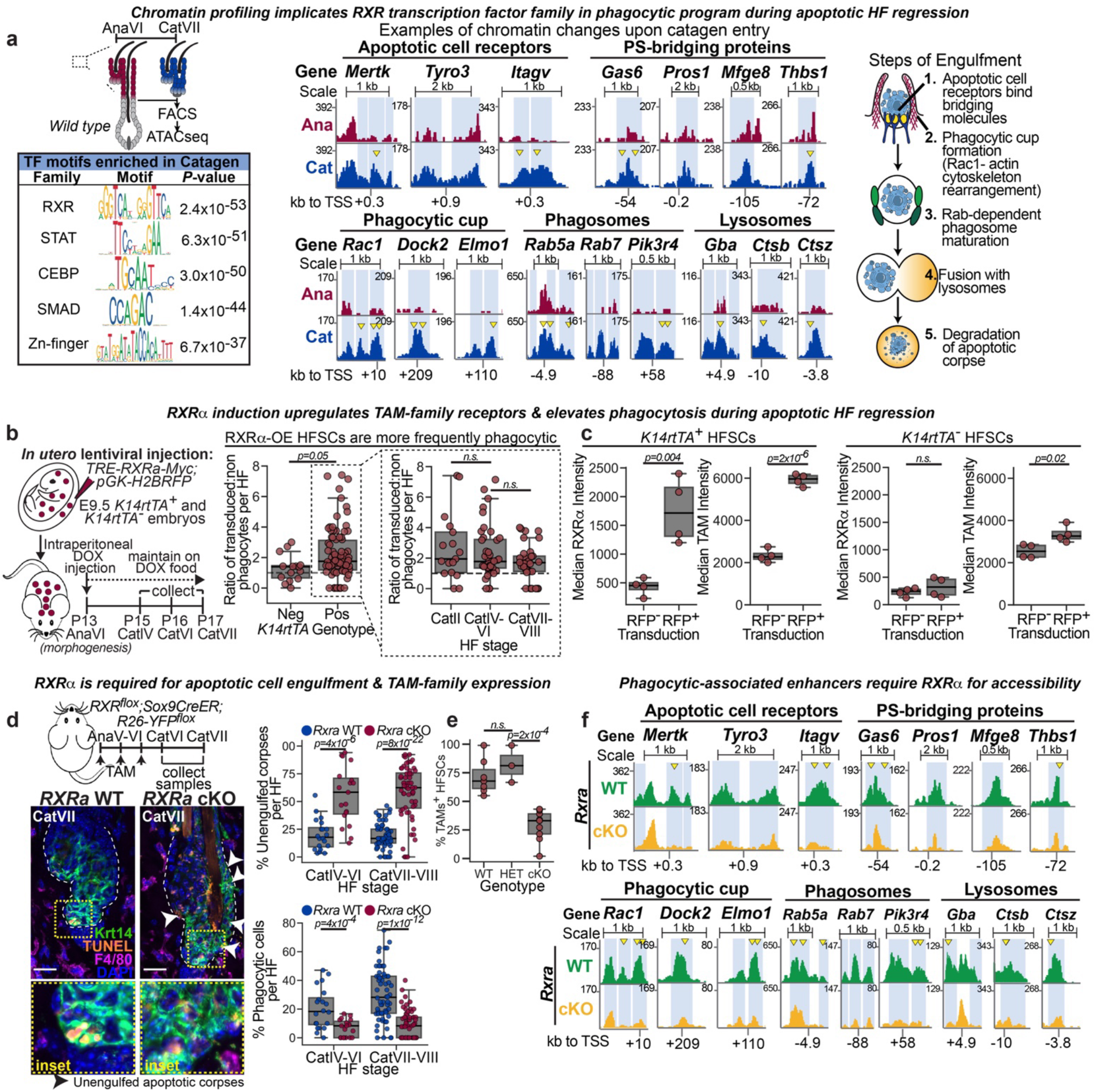
RXRa is a master regulator of the phagocytic hair follicle stem and progenitor cell state. **a,** Top left, Strategy to profile FACS-isolated hair follicle stem cells (HFSCs) in late anagen (AnaVI) versus late catagen (CatVII) by ATAC-seq. Bottom left, Motif enrichement analysis on catagen accessible peaks. Middle, Replicate-pooled peak tracks for enhancers associated with phagocytic genes in catagen (Cat) versus anagen (Ana). Transcription start site, TSS. Phagocytic genes are grouped according to engulfment step (schematized at right). Differential peaks highlighted in light blue, yellow arrowheads point to RXR-family bound footprints. Peak tracks in reads per genome coverage (RPGC). **b,** Left, Inducible RXRα expression strategy. Middle, Quantification of untransduced versus transduced phagocytic cells. n=2-4 mice per stage, 8-10 HF analyzed per mouse. **c,** Quantifications of RXRα (left) and TAM-family (right) expression in FACS-isolated HFSCs. Data are paired, with untransduced (RFP^-^) and RXRa-overexpressing (RFP^+^) cells co-occuring, n=4 mice. Data for *K14rtTA*^-^ control right-most panels. **d,** Left top, *Rxra* conditional knockout (cKO) strategy. Left bottom, Sagittal sections of late catagen *Sox9CreER;Rxra^fl/fl^;R26-YFP^fl/+^*skins. wild type, WT; Scale bar 20um. Right, Quantifications of unengulfed apoptotic bodies (top) and HFSCs containing corpses (bottom). 10-15 follicles analyzed per mouse, n=6-8 mice per genotype. **e,** Percentage of FACS-isolated TAM-family^+^ HFSCs per mouse for *Rxra* WT, heterozygous (HET) and cKO. n=3-8 mice per genotype. **f,** ATAC-seq pooled replicate peak tracks covering the enhancers in (a) for FACS-isolated CatVII HFSCs, scaled, normalized and annotated as in (a). All quantifications, multiple pairwise Student’s *T*-Tests, p-values indicated. n.s. not significant (p>0.05).

Recognition of corpses by phagocytic cells involves the activation of apoptotic cell clearance receptors, often by binding surface exposed phosphatidylserine (PS) on the apoptotic cell either directly or via the engagement of bridging molecules^4-6^ (**Fig. 2c**). To visualize the pattern of gene expression associated with apoptotic cell clearance receptors and bridging ligands, we created aggregate gene set scores for these terms and visualized them on the UMAP projected data. Strikingly, catagen HF stem and progenitor cells were strongly enriched for both apoptotic cell clearance receptors and bridging molecule expression, with a more modest increase in the lysosomal gene set (**Fig. 2d**, **Extended data Fig. 3b**). Among the genes differentially expressed in catagen were several members of the Tyro3/Axl/Mertk (TAM)-family receptor and integrin receptor pathways of apoptotic cell recognition and tethering, including the receptors *Tyro3*, *Mertk,* and *Itgav*, as well as the PS-bridging proteins *Gas6, Pros1, Mfge8* and *Thbs1* (**Fig 2d**, **Extended data Fig. 3c**, **Supplemental Table I**).

Turning to the physiological significance of our transcriptomic observations, we used FACS and observed a significant enrichment of TAM-family surface expression on HF stem and progenitor cells that had engulfed another cell (**Fig. 2e**, **Extended data Fig. 4a**). FACS quantifications documented that surface expression of TAM-family members peaked during catagen, but was largely absent at other phases of the hair cycle (**Extended data Fig. 4b,c**). A subset of these TAM-family^+^ HFSCs also displayed an expanded lysosomal compartment, indicative of apoptotic corpse degradation within phagolysosomes (**Extended data Fig. 4d**).

**Fig. 4.**
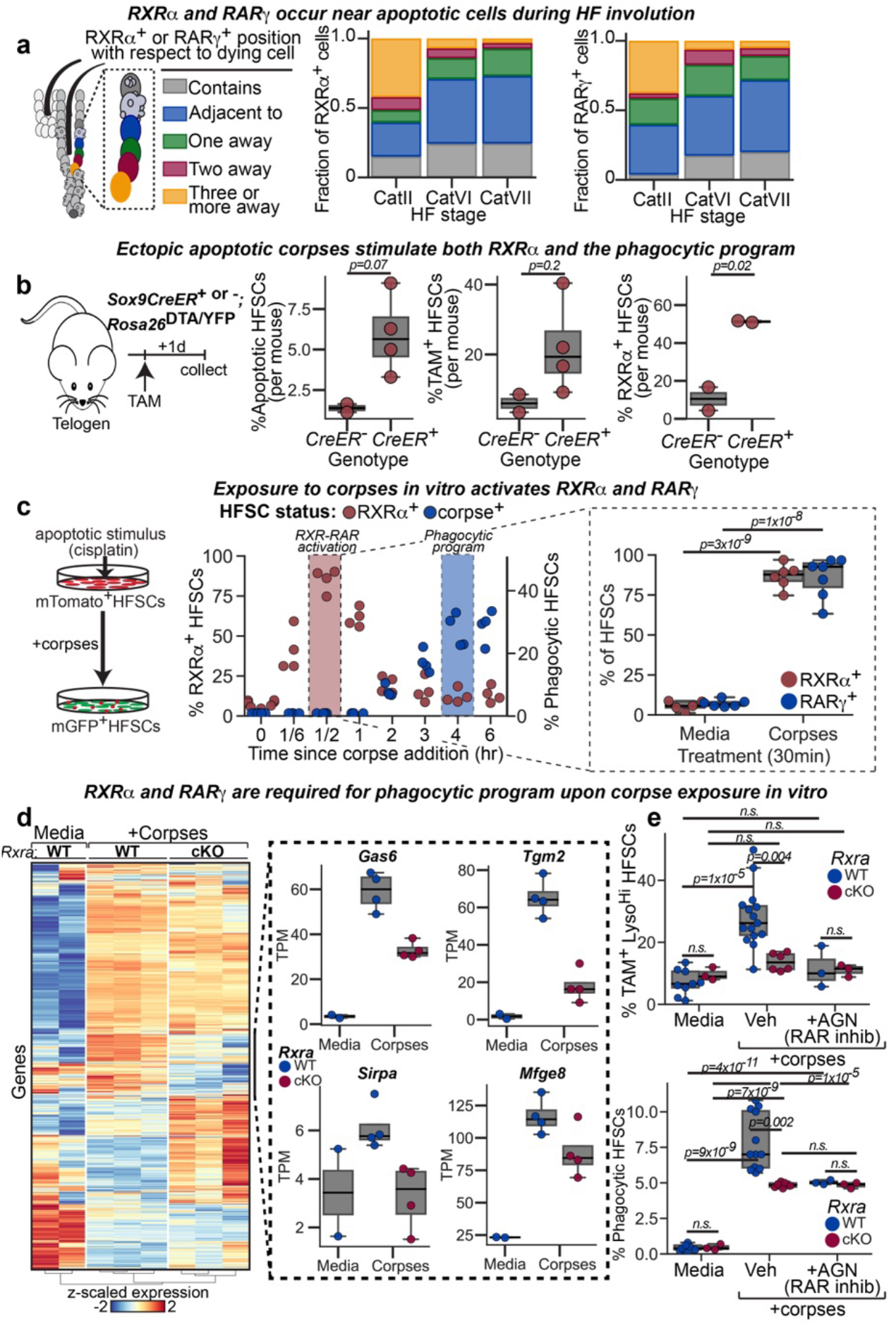
RXRα responds to corpse-derived signals to upregulate the phagocytic program. **a,** Left, Schematic of cell position with respect to nearest corpse. Fractions of RXRα^+^ (Middle) or RARγ^+^ (Left) position in relation to corpses across catagen. n= 300 cells across 3 mice per stage. **b,** Left, Induction of corpses in quiescent hair follicles (HFs). Right, Quantifications of total FACS-isolated HFSCs that are apoptotic (AnnexinV^+^), TAM-family^+^, and RXRα^+^ per mouse in control (*Sox9CreER^-^*) or corpse-positive (*Sox9CreER^+^*) animals. n= 2-4 animals per genotype. **c,** Left, *In vitro* strategy to expose naïve HFSCs to corpses directly. Middle, Time-course quantification of total HFSCs that are RXRα^+^ (red) versus containing a corpse (blue). Right, Quantification of RXRα^+^ and RARγ^+^ HFSCs at 30 min. post corpse addition. n=4-6 experiments per time point, performed in triplicate. **d,** Bulk RNA-squencing of *Rxra* wild type (WT) versus conditional knockout (cKO) HFSCs in media or upon corpse exposure. Left, Heatmap of differentially expressed genes (full list in Supplemental Table 4); Z-scaled expression scores of genes (rows; blue, downregulated; red, upregulated) by experimental replicates (columns; n=2-3 WT, 3 cKO). Right, Expression of phagocytic genes; TPM, transcript per million. **e,** Quantification of total HFSCs in culture that are both TAM-family^+^; Lysosome^high^ (top) or contain engulfed corpses (bottom) 4 hours post corpse addition to RXRα WT or cKO with or without RAR-family inhibition (+AGN condition). Data from n=3-8 experiments per genotype per condition, averaged across triplicates. All quantifications, multiple pairwise Student’s *T*-Tests, p-values indicated. n.s. not significant (p>0.05).

Interrogating the functional importance of our findings, we ablated *Mertk* in mice, and also blocked exposed phosphatidylserine with intradermally-injected AnnexinV. Both of these measures resulted in delayed apoptotic corpse clearance, with more unengulfed apoptotic cells and fewer phagocytic HF stem or progenitor cells compared to controls (**Fig. 2f,g**). Taken together, these results provide compelling evidence that HFSCs invoke a classical apoptotic cell clearance pathway, but do so transcriptionally and in a unique, highly controlled transient manner to act as periodic phagocytes.

### RXRa is a master regulator of the phagocytic state in hair follicle stem cells

The transient nature of the phagocytic program in HFSCs distinguished it from professional phagocytes whose primary role is to clear dead cells. To understand how the phagocytic program is dynamically regulated in tissue stem cells, we profiled the chromatin landscape of FACS-purified HFSCs at the end of the growth phase and during apoptotic regression. Using an assay for transposase-accessible chromatin by high-throughput sequencing (ATAC-seq) coupled with differential peak analysis, we identified two sets of dynamic chromatin peaks: those that closed upon entry to catagen, and those that became accessible (**Extended data Fig. 5**).

**Fig. 5.**
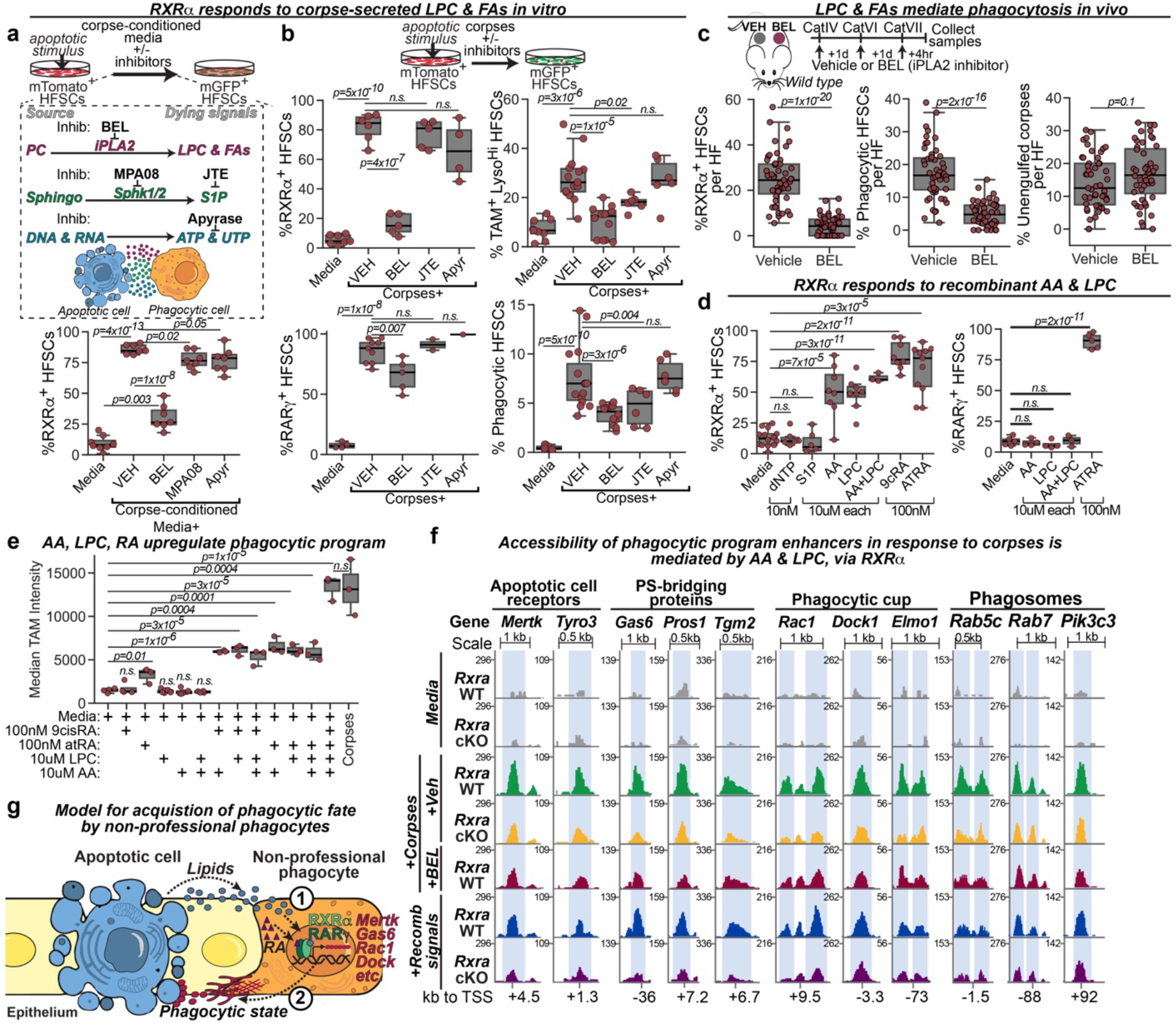
Corpse-derived lysophosphatidylcholine and fatty acids are necessary and sufficient for RXRα-mediated regulation of the phagocytic program. **a,** Top, Schematic of secreted signals from dying HFSCs; Inhib, inhibitors; BEL, bromoenol lactone; PC, phosphatidylcholine; LPC, lysophosphatidylcholine; FAs, fatty acids; JTE, JTE013; Sphingo, sphingosine; S1P, sphingosine-1-phosphate. Bottom, Quantification of RXRα^+^ HFSCs responding to corpse-conditioned media. Apyr, Apyrase; n=7 experiments per condition **b,** Top, Experimental schematic. Left, Percentages of RXRα^+^ (Middle) and RARγ^+^ (Bottom) HFSCs per experiment, 30 min. post corpse addition. n= 4-6 experiments per condition, as averaged triplicates. Right, Percentages of TAM-family^+^; Lysosome^high^ HFSCs (Middle) and corpse-containing HFSCs (Bottom) per experiment, 4 hours post corpse addition. n= 6-12 experiments per condition. **c,** Top, Strategy to inhibit production of LPC and FAs during catagen by intradermal injections of the iPLA2 inhibitor BEL in WT mice. Contralateral vehicle injections serve as control, n= 6 mice. Bottom, Percentages of RXRα^+^ cells (Left), corpse-containing HFSCs (Middle), and unengulfed apoptotic bodies (Right) per HF. n=6-8 follicles quantified per condition per mouse, n=6 mice. **d,** Percentage of RXRα^+^ (Left) and RARγ^+^ (Right) HFSCs per experiment, 30 min after addition of the indicated recombinant molecules; dNTP, 10nM each dUTP and dATP; AA, arachidonic acid; 9cRA, 9-*cis* retinoic acid; ATRA, all-*trans* retinoic acid. n=4-12 experiments per condition, as averaged triplicates. **e,** FACS quantification of TAM-family expression on HFSCs treated with the indicated recombinant molecules compared to TAM expression post corpse exposure, all 4 hours later. n=3-6 experiments per condition. **f** ATAC-seq replicate-pooled peak tracks for enhancers associated with phagocytic genes in cultured *Rxra* WT and cKO HFSCs exposed to apoptotic corpses with (+Veh) or without (+BEL) secreted LPC and FAs, or treated with retinoic acid+LPC+AA (+Recomb signals) for 4 hours. Peaks with differential accessibility highlighted in light blue. Peak tracks in reads per genome coverage (RPGC). Lysosome-associated peaks in Extended Data Fig. 10. **h,** Model by which HFSCs sense apoptotic corpses via secreted LPC and FAs, and activate RXR-RAR signaling to adopt a transient phagocytic fate. All quantifications, multiple pairwise Student’s *T*-Tests, p-values indicated. n.s. not significant (p>0.1).

Of note, genes in proximity to catagen-opening peaks encoded proteins associated with apoptotic cell receptors, soluble PS-bridging molecules, phagocytic cup formation and phagolysosome maturation (**Fig. 3a**). To identify candidate transcription factors mediating this dynamic chromatin accessibility, we performed motif enrichment analysis for peaks that either opened or closed during the regressive phase. We discovered that motifs belonging to the retinoid X receptor (RXR)-family of transcription factors were significantly enriched in peaks gained in catagen (**Fig. 3a**). Notably, the most enriched RXR-family motif among catagen-specific peaks was a direct-repeat 2 (DR2) motif composed of hexameric RXR binding motifs separated by 2 nucleotides. This repeat has been implicated in RXR heterodimerization with retinoic acid receptors (RARs)^16-18^ (**Extended Data Fig. 6a**).

To test the hypothesis that RXR-RAR family members regulate the phagocytic machinery, we first surveyed the expression of the entire nuclear receptor superfamily across the hair cycle. Of these, *Rxra* and *Rarg* stood out as having appreciable expression that peaked at the height of apoptotic regression in the HF (**Extended Data Fig. 6b**). Immunofluorescence confirmed the nuclear localization of both RARγ and RXRα in the proximity of dying cells in catagen (**Extended Data Fig. 6c,d**).

If RXRα-RARγ is the signaling complex responsible for the dynamic regulation of the catagen-triggered phagocytic program, changing the levels of RXRα (the obligate heterodimeric partner) should impact this step. To do so, we first ectopically expressed RXRα in skin progenitors. Using our powerful ultrasound-guided *in utero* lentiviral delivery method, we transduced skin progenitors of *K14rtTA* embryos with a constitutively expressed RFP transduction control gene and a doxycycline-inducible human *RXRA* transgene. We then administered doxycycline at the end of the morphogenesis growth phase of the first hair cycle and followed the fate of *RXRA* super-activated HF stem and progenitor cells during catagen. Congruent with a role for RXRα signaling in induction and/or maintenance of this transient phagocytic phase, catagen ORS cells with higher RXRα more frequently contained engulfed apoptotic bodies than neighbouring untransduced cells (**Fig. 3b**, **Extended Data Fig. 7a**). Relative to controls, HFSCs with elevated RXRα also displayed significantly higher levels of TAM-family receptors, consistent with enhanced phagocytic ability (**Fig. 3c**).

We next turned to evaluating the functional consequences of eliminating RXRα from catagen-phase HFSCs. For this, we bred our tamoxifen-inducible *Sox9CreER;R26-YFP-floxed* reporter mice to *Rxra-floxed* animals, administered tamoxifen at the end of anagen, and then assessed HFSC phagocytic ability during the subsequent destructive phase. In the absence of RXRα, catagen HFs initiated apoptosis normally, but were strikingly defective at clearing corpses (**Fig. 3d, Extended Data Fig. 7b,c**). Notably, significantly fewer HF stem and progenitor cells contained engulfed corpses and correspondingly, interstitial spaces were littered with apoptotic debris. This in turn attracted F4/80-positive macrophages, which contacted *Rxra* conditional knockout HFs (**Fig. 3d**).

The appearance of macrophages suggested that the catagen HFSCs stopped functioning as non-professional phagocytes in the absence of RXRα. Indeed, RXRα-deficient HFSCs failed to upregulate TAM-family receptor expression in late catagen (**Fig. 3e**). To assess the ability of RXRα to regulate phagocytic receptor expression directly, we mosaically transduced skin progenitors of *Sox9CreER; R26-LSL-Cas9-EGFP* embryos with a lentivirus harboring an *Rxra-*targeting sgRNA and a mScarlet reporter, and then administered tamoxifen at the end of the first anagen. By catagen, Scarlet^+^GFP^+^ cells, which had received both sgRNA and activated Cas9, were largely deficient for RXRα protein and lacked TAM-receptors, whereas single positive cells containing either active Cas9 or sgRNA maintained high levels of both RXRα and surface TAM-family receptors (**Extended Data Fig. 7c**). Together, these data provided compelling evidence that RXRα is required and sufficient to cell autonomously activate a phagocytic program in catagen-phase HFSCs to clear apoptotic neighbours.

Given RXRα’s role as a nuclear receptor, we next addressed whether RXRα participated in transcriptional regulation of apoptotic cell clearance. To do so, we first assessed whether RXRα is functionally important for opening the dynamic enhancer peaks that emerge during the transition from anagen to catagen phases of the hair cycle. ATAC-seq and differential peak analyses of FACS-isolated late catagen-phase HFSCs revealed that 8,311 peaks altered chromatin accessibility when *Rxra* was ablated, while 4,222 peaks of these required RXRα for accessibility (**Extended Data Fig. 7d-h**). Notably, the apoptotic cell clearance enhancer peaks were among the RXRα-dependent peaks which gained accessibility during catagen (**Fig. 3f**).

Our data thus far suggested that RXRα specifically regulates the phagocytic transcriptome of HF stem and progenitor cells, which should therefore be perturbed upon transcriptional antagonism of RXR-family members. To test this hypothesis, we transiently administered the RXR antagonist HX531^18,19^ during catagen. In contrast to the vehicle control (injected into contralateral back skin), RXR-inhibition decreased the number of phagocytic HFSCs and increased the number of unengulfed apoptotic corpses in late catagen (**Extended Data Fig. 7i,j**). Similar to *Rxra* cKO animals, transcriptional antagonism of RXR led to decreased TAM-family and *Mfge8* expression *in vivo*.

Together with expression patterns, these functional data strongly implicated RXRα-RARγ in the direct regulation of a suite of apoptotic cell clearance genes necessary to transiently adopt a phagocytic state in HF stem and progenitor cells. In this regard, it was intriguing that activities of RXRα with different nuclear receptors have been reported to enhance the phagocytic program in macrophages^20-23^. Our findings highlight an unexpected convergence between professional and non-professional phagocytes in how apoptotic cell clearance is achieved at the transcriptional level, and suggest that by binding to different partners, RXRs may fine-tune the regulation of this process to suit the needs of different tissues.

### RXRα-RARγ complex senses nearby corpses

Efficient apoptotic cell clearance by non-professional phagocytes is still poorly understood. Early embryonic epithelia appear to shuttle apoptotic corpses long distances to reach phagocytic trophoblast cells^24^. By contrast, live imaging has indicated that embryonic and adult skin epithelia clear apoptotic corpses that are in close proximity^13, 25^. Such observations suggest that spatially constrained epithelial cells harbor the ability to efficiently sense, respond and clear neighboring apoptotic cells to deal with stochastic cell death.

To begin to identify the underlying mechanisms involved, we first examined the spatial relation between apoptotic corpses and RXRα-RARγ during HF regression. Throughout catagen, many RXRα- and RARγ-positive cells were found within one cell body distance of an unengulfed corpse (**Fig. 4a**, **Extended Data Fig. 8a**). Probing this relation more deeply, we next ectopically induced apoptosis in telogen HFs by sparsely activating diphtheria toxin subunit A (DTA) in a subset of HFSCs during the resting phase of the hair cycle. This resulted in a dramatic rise in RXRα^+^, TAM-family receptor^+^ expression within the healthy (DTA^neg^) HF stem and progenitor population (**Fig. 4b, Extended Data Fig. 8b**). Intriguingly, forced overexpression of RXRα in the absence of apoptotic cells was not sufficient on its own to alter phagocytic receptor expression from baseline (**Extended Data Fig. 8c**). Taken together, these results indicated apoptotic corpses are an essential trigger for the activation of RXRα and the upregulation of TAM-family receptors *in vivo*.

Given the close spatial relation between activation of RXRα-RARγ in HFSCs and apoptotic corpses *in vivo*, we posited that HFSCs may be able to directly sense corpses without input from other cell types. To test this, we cultured naïve telogen HFSCs and exposed them to apoptotic corpses. Within 30 minutes of corpse addition, nuclear RXRα and RARγ were strongly elevated, followed by apoptotic cell engulfment plateauing around 4-6 hours later (**Fig. 4c, Extended Data Fig. 8d**). Pre-treatment of apoptotic corpses with either recombinant annexinV (to mask exposed phosphatidylserine) or BMS-777607 (to inhibit TAM-family receptor activity) impaired their HFSC-stimulated engulfment, suggesting that a similar phagocytic program was operative *in vitro* as *in vivo* (**Extended Data Fig. 8e**). Indeed, in response to corpses, cultured HFSCs upregulated TAM-family receptor surface expression and engulfed apoptotic bodies both of which were blocked by transcriptional antagonists against either RAR or RXR families, further consistent with the notion that the phagocytic program is dependent upon apoptotic corpses and driven by RARγ-RXRα activities (**Extended Data Fig. 8f**).

To directly assess the requirement for RXRα in mediating the response to corpses *in vitro*, we cultured FACS-isolated YFP+ telogen-phase HFSCs from *Rxra* WT and cKO mice and transcriptionally profiled them 4 hours after corpse addition. WT HFSCs responded to corpse exposure by transcriptionally upregulating a cohort of phagocytic genes (**Supp Table for full list**). Notably, a subset of these genes showed a markedly diminished response in *Rxra* null HFSCs concomitant with functionally impaired apoptotic corpse clearance (**Fig. 4d,e**). Finally, although significantly affecting apoptotic corpse clearance on its own, the RAR-family inhibitor AGN 193109 had no further effect on *Rxra* null cells, suggesting that the two act cooperatively rather than in parallel (**Fig. 4e**, **Extended Data Fig. 8g**). Altogether, our culture data demonstrated that HFSCs directly sense the presence of corpses and respond by upregulating a phagocytic state that requires RXRα-RARγ signaling.

### Hair follicle stem cells detect corpses via secreted signals to activate RXR-RAR activity

Macrophage elevation of TAM-family receptors via RXR-PPAR/LXR signaling requires engulfment and digestion of corpses^20-23^. By contrast, in HFSCs neither engulfment nor digestion of corpses was required for nuclear RXRα/RARγ or downstream TAM-family and lysosomal expression (**Extended Data Fig. 9a**). These data suggest that the mechanism of corpse activation of RXR signaling may differ in non-professional versus professional phagocytes and depends on both distinct co-receptors and ligands.

Interestingly, when we titrated apoptotic corpses to <1 per 100 HFSCs *in vitro*, we saw local increases in RXRα/RARγ-positive cells around each corpse (**Extended Data Fig. 9b**), reminiscent of that seen throughout catagen *in vivo*. That said, corpse-conditioned media increased the percentage of RXRα-positive cells to the same levels as corpses themselves, indicating that a secreted corpse factor(s) is signaling their presence (**Fig. 5a,b**). The best characterized “find-me” signals secreted by apoptotic cells are free nucleotides, sphingosine-1-phosphate (S1P) and lysophosphatidylcholine (LPC)^25-30^. Using small molecule inhibitors to block the generation of S1P (by inhibiting Sphk1/2 with MPA08), LPC (by inhibiting iPLA2 mediated phosphatidylcholine cleavage with bromoenol lactone, BEL) or free nucleotides (by inducing their degeneration with recombinant apyrase, Apyr), we found that of these three, LPC was the only one required to activate RXRα-RARγ (**Fig. 5a,b**). Correspondingly, blocking phosphatidylcholine cleavage by apoptotic corpses abolished both TAM and lysosome upregulation in HFSCs as well as corpse engulfment (**Fig. 5b**).

RNA sequencing verified that the effects of impairing phosphatidylcholine cleavage were at the transcriptional level, with moderate downregulation in a subset of apoptotic cell clearance receptors, PS-bridging proteins, actin cytoskeletal regulators involved in phagocytic cup formation, and mediators of phagolysosome maturation (**Extended Data Fig. 9c**). Finally, eliminating phosphatidylcholine cleavage/LPC production during catagen *in vivo* significantly diminished both RXRα upregulation in HF stem and progenitor cells adjacent to corpses, and also the number of corpse-containing HFSCs (**Fig. 5c**). By contrast, blocking the generation of S1P, free nucleotides, or exposure of phosphatidylserine *in vivo* showed no overt effect on RXRα activation or the phagocytic program (**Extended Data Fig. 9d**). Together, our data point to the view that in apoptotic cells, the cleavage of phosphatidylcholine to generate LPC and free fatty acids (FAs)^26,31^ acts locally to activate RXR signaling and induce a phagocytic state in neighbouring HFSCs. Given that blood LPC levels are much higher than those derived from dying cells *in vivo*, a role for apoptotic cell-derived LPC in stimulating tissue phagocytes resolves the conundrum regarding its biological activity as a chemotactic molecule for circulating professional phagocytes ^4,26,32-34^.

Caspase-mediated activation of iPLA2 leads to hydrolysis of phosphatidylcholines at the sn-2 position, and subsequent release of LPC and free FAs^26,31,35^. A major constituent at the sn2 position in mammalian cell membrane phospholipids is the fatty acid arachidonic acid (AA)^36,37^, which has been described as a natural ligand for RXR^38,39^. In this regard, adding recombinant AA and/or LPC elevated nuclear RXRα intensity in roughly 50% of cultured HFSCs across a physiologically relevant range of concentrations^26,40^ (**Fig. 5d, Extended Data Fig 9e,f**).

In contrast to RXRα, RARγ did not respond to any concentration or combination of AA or LPC tested, consistent with its classic ligands being 9-*cis* retinoic acid (9cRA)^41,42^ and all-*trans* retinoic acid (ATRA)^43,44^. Notably, however, the combination of retinoic acid (RA) isomer and LPC (and/or AA) induced high levels of both nuclear RXRα and RARγ in nearly all HFSCs *in vitro* (**Extended Data Fig. 9f**). Additionally, TAM receptor expression comparable to corpse-exposure could be achieved by the combination of AA, LPC and both RA isoforms (**Fig.5e**, **Extended Data Fig. 9g**). Optimal activation of the transcriptional phagocytic program was also seen as judged by elevated transcripts for the TAM genes *Tyro3, Axl,* and *Mertk* and the PS-bridging genes *Gas6, Mfge8 and Thbs1* (**Extended Data Fig. 9h**).

To unequivocally document and characterize the dependency of phagocytic program genes on RXRα-RARγ activation, we used ATAC-seq and profiled WT and *Rxra* cKO HFSCs in two complementary settings: 1) in response to corpses with (+VEH) or without (+BEL) phosphatidylcholine hydrolysis/LPC+AA generation, and 2) in response to AA, LPC and RA combined. In the absence of corpses or signaling cocktail, phagocytic genes were in a closed chromatin state (**Fig. 5f, Extended Data Fig. 10a-e**). Upon corpse exposure, 12,614 peaks lost while 5,004 peaks gained accessibility. Approximately half of the peaks gaining accessibility were markedly diminished upon *Rxra* ablation. These peaks were also sensitive to the presence of LPC/FAs as documented by their decline upon exposure to BEL-treated corpses. Many of these peaks resided within putative enhancers for genes involved in multiple stages of apoptotic cell clearance *in vivo* as well as *in vitro* (compare **Figs. 3f** and **5f**). Impressively, the cocktail of AA, LPC and RA recapitulated the effect of corpses on approximately one third of the RXRα-dependent peaks including those in enhancers of genes encoding apoptotic cell recognition, engulfment, and processing pathways (**Fig. 5f, Extended Data Fig. 10f**).

In summary, our findings reveal that tissue stem and progenitor cells can be stimulated by local apoptotic corpse-derived factors to activate a phagocytic program that efficiently and expeditiously rids the tissue of dying cells that might otherwise trigger an inflammatory response (**Fig. 5g**). We discovered that in contrast to professional phagocytes, tissue stem cells fine-tune this program and are only diverted from their normal tasks when they receive the right stimulatory signals. For the HF, these signals appear only transiently in the destructive phase of the hair cycle when TGFβ causes cells within the HF to either terminally differentiate (the hair shaft) or apoptose (the ORS)^13,45^. The stem cells maintain tissue homeostasis on multiple fronts through a mechanism that is rooted in their ability to sense apoptotic corpse signals, primarily the lipids LPC and AA, in combination with local tissue signals, e.g. retinoic acid. In combination, these factors induce nuclear RXRα-RARγ signaling to orchestrate a transient phagocytic state in the epithelial stem cells, which enables them to efficiently engulf and clear apoptotic cells in a highly localized and temporally controlled manner.

We surmise that this method of sensing apoptotic corpses may be broadly applicable to non-professional phagocytes in other epithelial tissues that have to balance sporadic cell death with maintaining an intact barrier. Indeed, many of these tissues utilize the same phagocytic receptors and PS-bridging molecules that converge on the general Rac1-Elmo-Dock pathway that mediates phagocytic cup formation and subsequent phagolysosome maturation^46-53^. That said, the beauty of having the pathway dependent upon RXRs is that while corpse-dependent production of lyso-lipids and free fatty acids is the universal activator of this transcription factor, RXRs can heterodimerize with a diverse family of binding partners each of which have their own specific small molecule ligands, differentially produced under different tissue conditions. By having a combinatorial trigger, phagocytosis can be tailored to suit the specific needs of each tissue. Additionally, the ability of stem and progenitor cells to serve as major contributors to apoptotic cell clearance provides a powerful mechanism for maintaining tissue homeostasis while enabling stem cells to reside in immune-privileged niches devoid of professional phagocytes.^46,54^ The transient phagocytic state we describe here, which is tuned specifically to the presence of nearby apoptotic corpses, offers an attractive mechanism to balance phagocytic duties against the primary stem cell function of replenishing differentiated cells to maintain tissue fitness.

---

**Extended Data Fig. 1.**
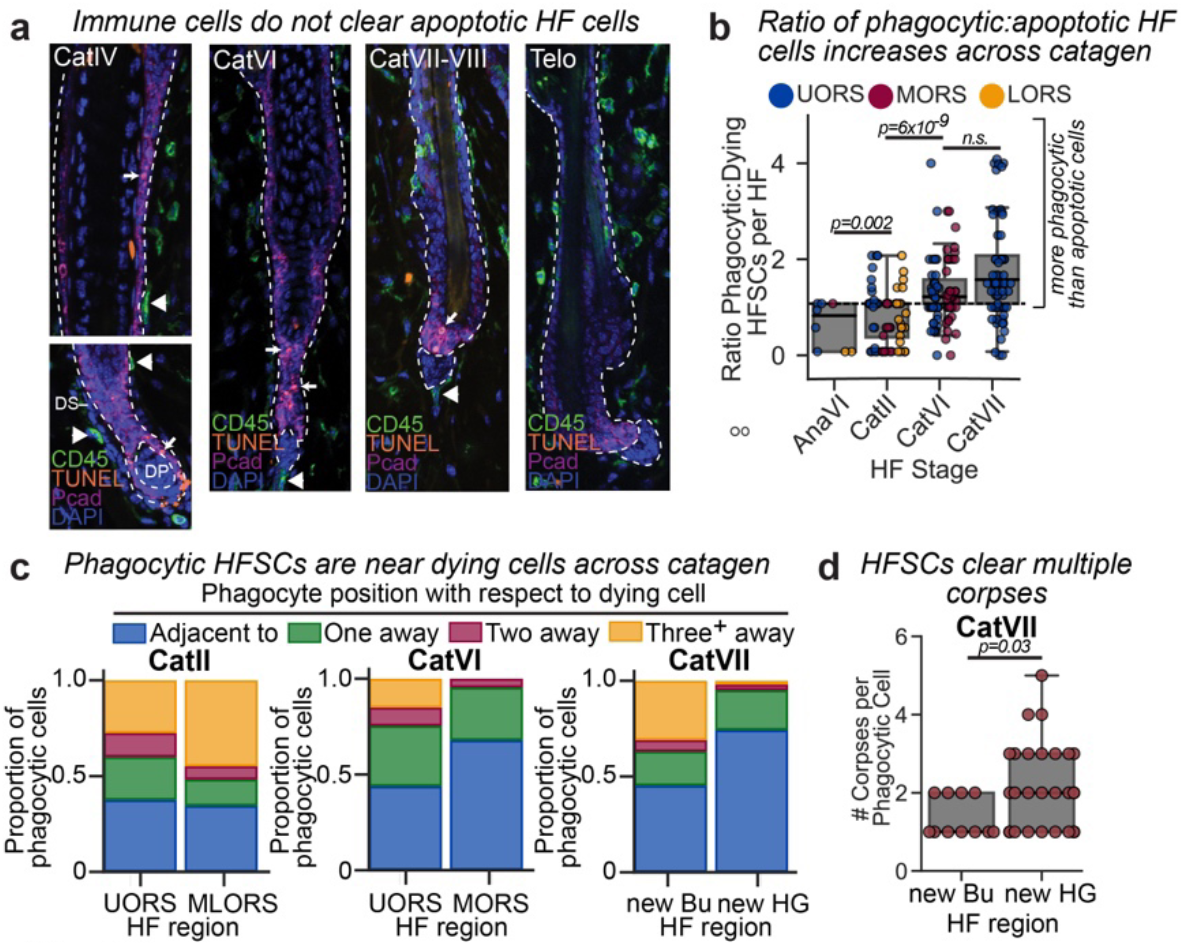
*(Related to Fig.1)* Apoptotic corpses are cleared by neighbouring hair follicle stem cells in catagen. **a,** Sagittal skin sections of CD45^+^ immune cells in the dermis (white arrowheads) that do not contact TUNEL^+^ apoptotic corpses (white arrows) within the regressing hair follicle, outlined by P-cadherin (Pcad, purple). Cat, catagen stage; Telo, telogen. Images are representative of 25-30 follicles across 3 mice per stage. Dashed lines denote dermo-epithelial border. **b,** Ratio of phagocytic to apoptotic cells per hair follicle (HF) across the hair cycle. 10-20 follicles were quantified per mouse, *n*=4-6 mice per stage. **c,** Quantification of HF phagocytic cell position with respect to nearest apoptotic cell, broken down by position within the outer root sheath (ORS). UORS, upper ORS; MLORS, middle and lower ORS; MORS, middle ORS; Bu, bulge; HG, hair germ. **d,** Quantification of the number of apoptotic corpses contained per phagocytic stem (new Bu, new bulge) and progenitor (new HG, new hair germ) cell via transmission electron micrography. *n*=7 hair follicles from 2 wild type mice. All quantifications, multiple pairwise Student’s *T*-Tests, p-values indicated. n.s. not significant (p>0.05).

**Extended Data Fig. 2.**
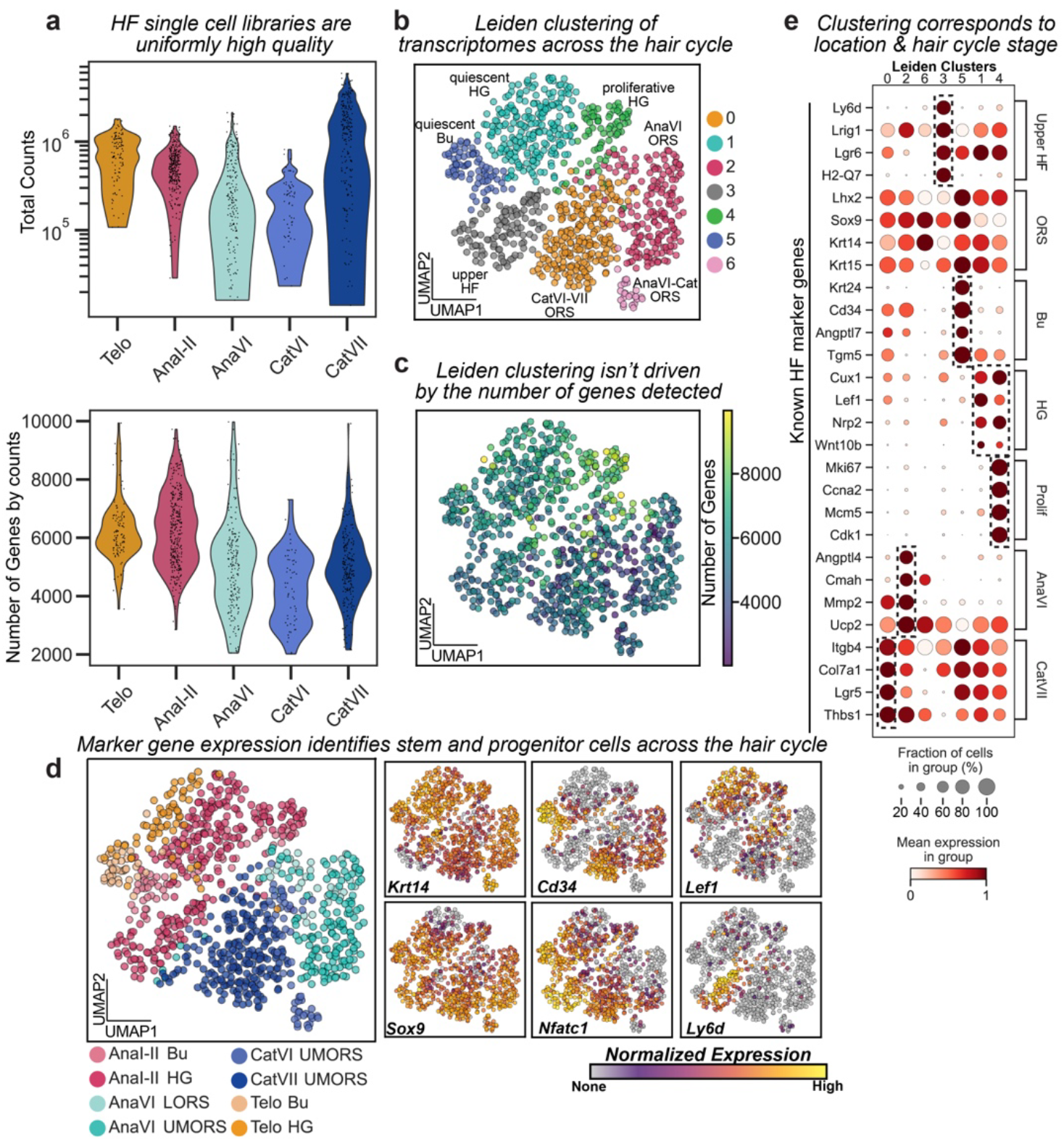
*(Related to Fig. 2)* Single-cell RNA-sequencing of hair follicle epithelial cells across the hair cycle. **a,** Total counts per cell (Top) and number of genes detected per cell (Bottom) for single-cell transcriptomic libraries across the hair cycle. Telo, telogen; AnaI-II, early anagen; AnaVI, late anagen; CatVI-VII, mid-late catagen. **b,** Uniform manifold projection (UMAP) representation of single cell transcriptomes coloured according to Leiden-based clustering algorithm. Cluster colour legend is at right, putative cluster identity based on hair follicle (HF) cycle stage and marker gene expression annotated on UMAP. **c,** UMAP representation of the number of genes detected per cell (colourbar legend at right). **d,** Left, UMAP representation coloured by anatomic location (via FACS antibody markers) and hair cycle stage. Bu, bulge; HG, hair germ; LORS, lower outer root sheath (ORS); UMORS, upper-middle ORS. Right, UMAPs of relative expression levels (log2[TPM+1]) of selected marker genes. ORS markers: *Krt14, Sox9*; Bu hair follicle stem cell (HFSC) markers: *Cd34, Nfatc1*; HG progenitor marker: *Lef1*; Upper HF marker: *Ly6d*. **e,** Dot plot representation of expanded marker gene analysis across Leiden clusters, with each vertical column corresponding to one grouped Leiden cluster and each horizontal row representing a marker gene. Marker genes are organized according to their reported function and location with the hair cycle; Prolif, proliferation. Dot size corresponds to the fraction of cells expressing each gene. Dot colour corresponds to normalized mean expression level within each cluster.

**Extended Data Fig. 3.**
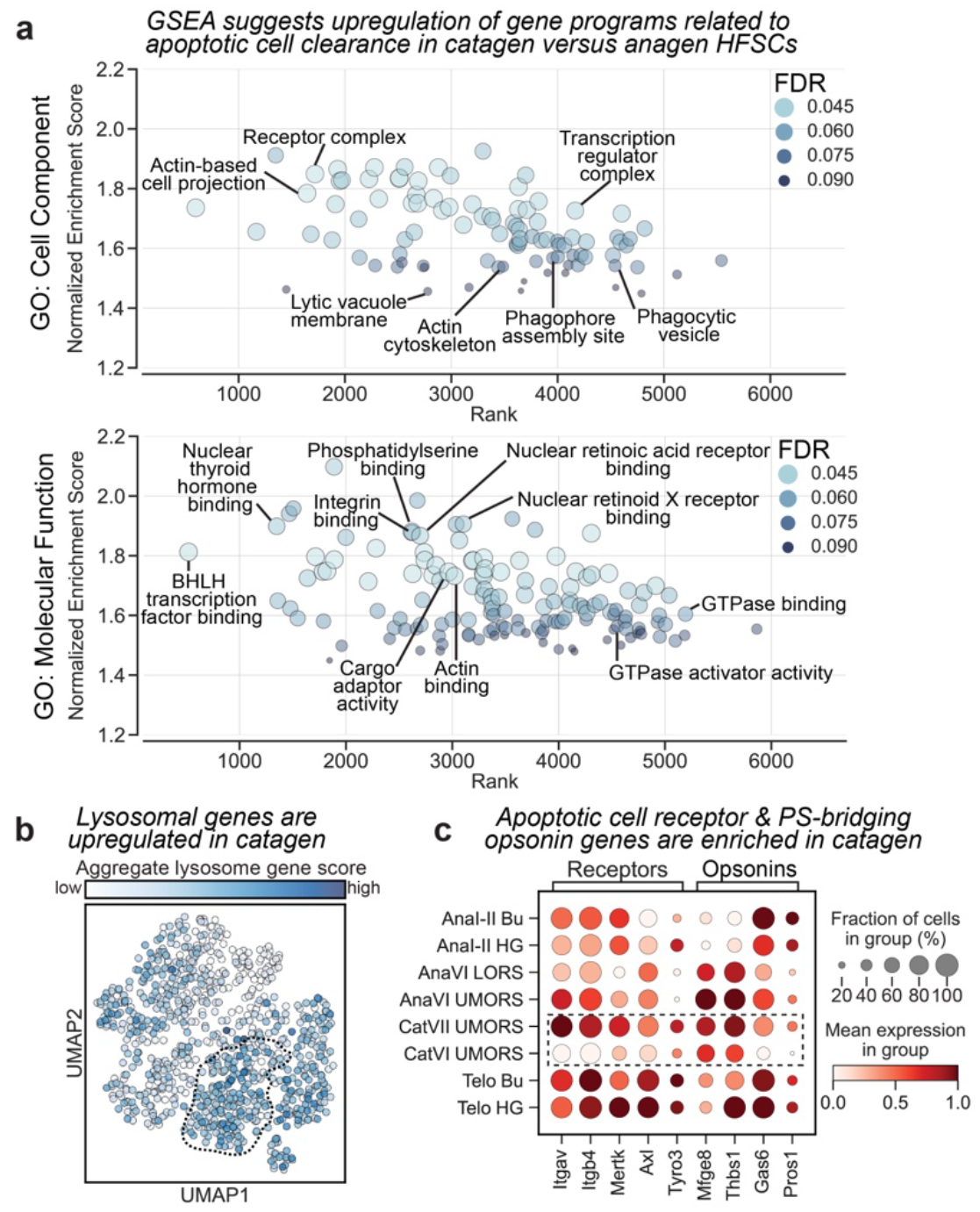
*(Related to Fig. 2)* Gene set enrichment identifies pathways related to apoptotic cell clearance upregulated in catagen hair follicle stem and progenitor cells. **a,** Visual representation of all gene sets identified as upregulated in catagen versus anagen (FDR <0.1), broken by Gene Ontology (GO) category. Gene set enrichment analysis (GSEA) ranks gene set terms by position at which maximum enrichment occurs (horizontal axis), and by normalized enrichment score (vertical axis). Dot colour and size reflects the False Discovery Rate (FDR) calculated for each gene set (legend inset). Selected gene sets related to apoptotic cell clearance are annotated. See Supplementary Tables 1 and 2 for full lists. **b,** UMAP representation of single-cell transcriptomes coloured by aggregate lysosomal gene set score shows enrichment in the catagen cluster (dashed black outline). **c,** Dot plot representation of selected apoptotic cell recognition receptors and phosphatidylserine-bridging genes (vertical columns) across hair cycle stages (horizontal rows). Each dot represents the fraction of cells (size) and normalized mean expression (colour) of a single gene across the hair cycle grouped single cell transcriptomes.

**Extended Data Fig. 4.**
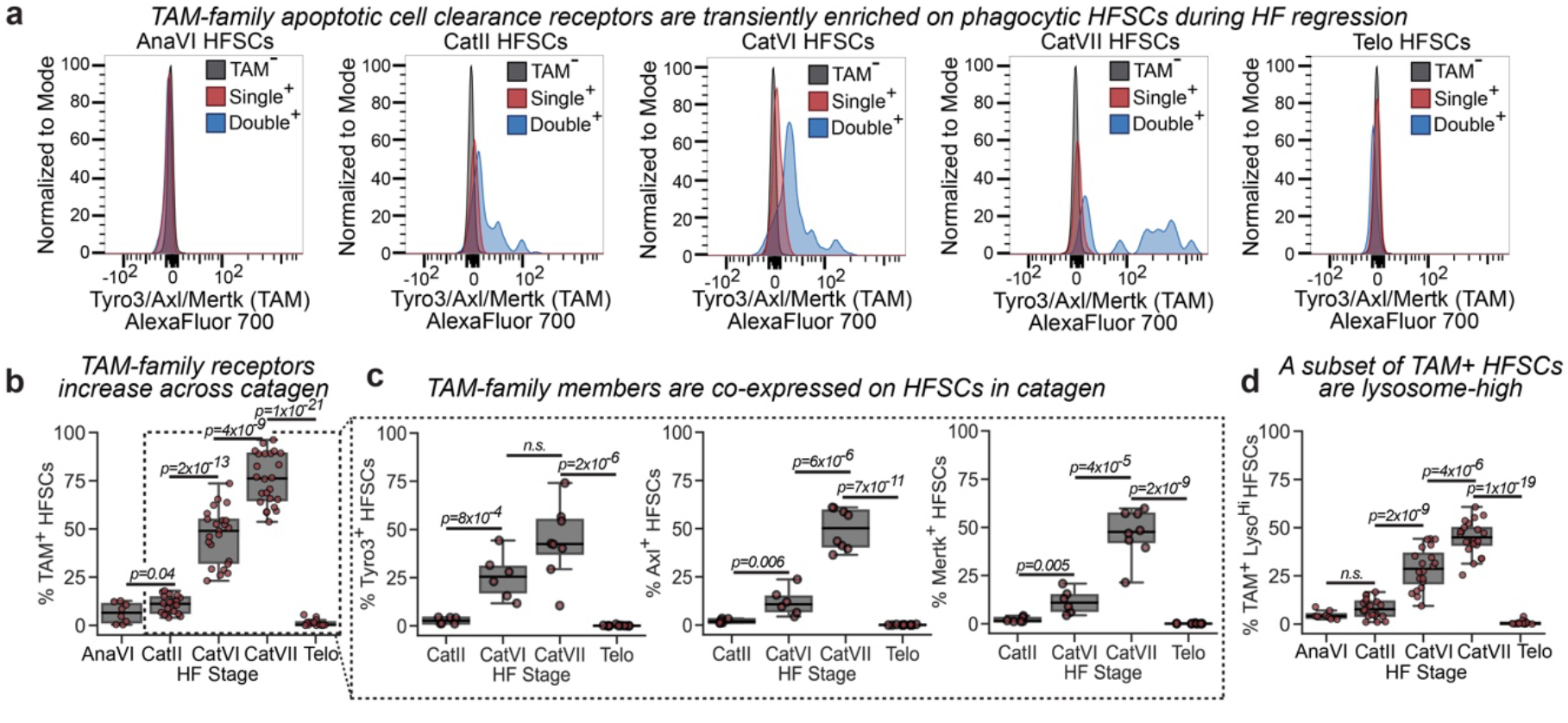
*(Related to Fig. 2)* TAM-family receptor pathway for apoptotic cell engulfment is expressed in hair follicle stem and progenitor cells during apoptotic regression. **a,** FACS plots showing normalized counts of cell-surface expression of *Tyro3/Axl/Mertk* (TAM)-family receptors on hair follicle stem cells (HFSCs) across the hair cycle, from the end of anagen growth phase (AnaVI), through early (CatII), mid (CatVI) and late catagen (CatVII) to quiescence (Telo, telogen). *Sox9CreER^+^* was used to drive *Brainbow2.1* fluorophore reporter expression so that all HFSCs are stochastically labeled with a single fluorophore (single+) and only phagocytic HFSCs (containing an apoptotic corpse of a different colour) can be double+. See supplemental methods for FACS sorting scheme. *n*= 4-6 mice analyzed per stage. Data is quantified in Fig. 2d. **b,** Percentage of TAM-family^+^ HFSCs per mouse across the hair cycle. *n*=8-16 mice per hair follicle (HF) stage. **c,** Percentage of FACS-isolated HFSCs expressing each individual TAM-family member (Tyro3, left; Axl, middle; Mertk, right) during catagen. *n*= 6-10 mice analyzed per stage. **d,** Percentage of HFSCs which are TAM^+^ and lysosome^high^ across the hair cycle. *n*= 8-16 mice per stage. All quantifications, multiple pairwise Student’s *T*-Tests, p-values indicated. n.s. not significant (p>0.05).

**Extended Data Fig. 5.**
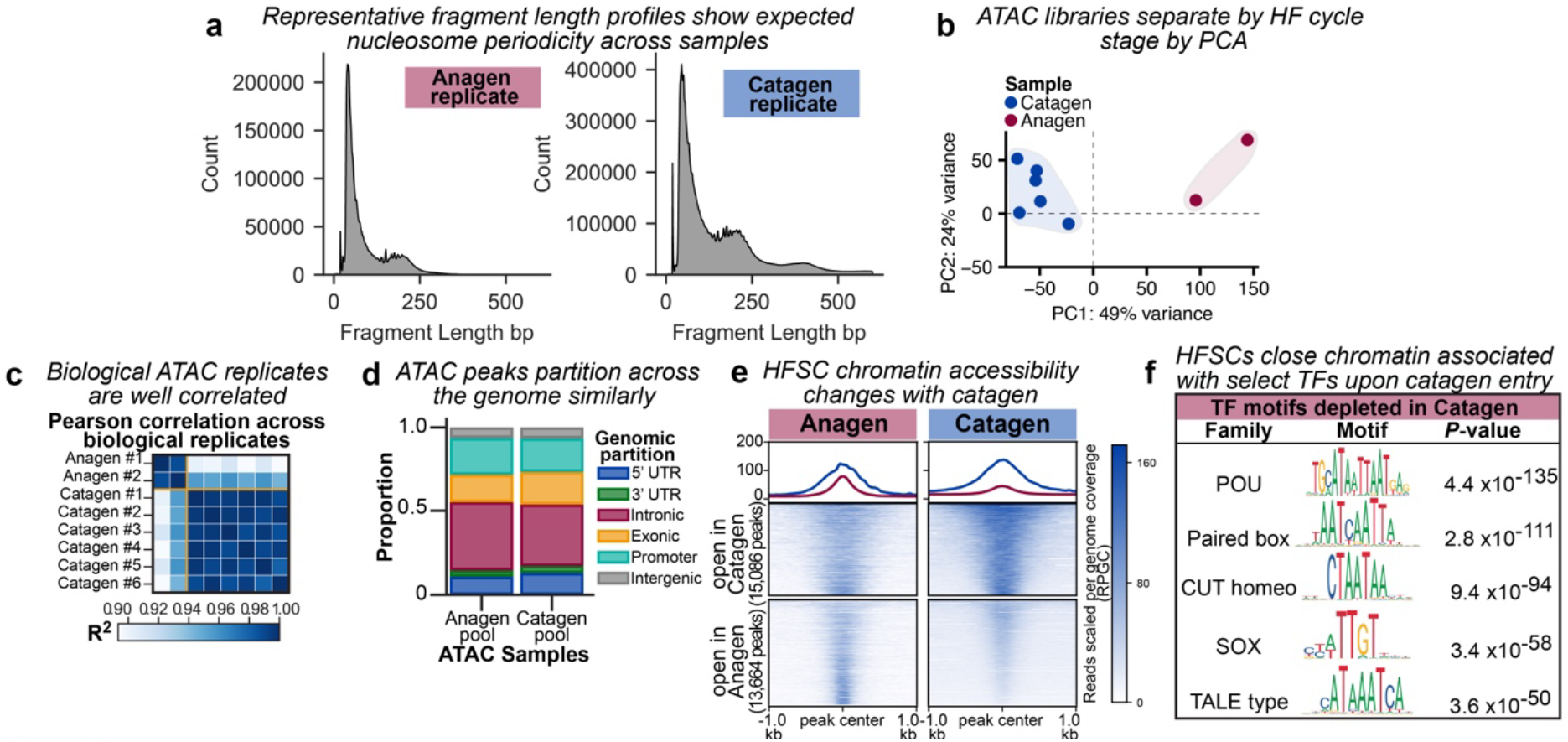
*(Related to Fig. 3)* Profiling hair follicle stem cells from growth to regression reveals a shift in chromatin accessibility associated with apoptotic elimination. **a,** Representative histograms of fragment lengths in base pairs (bp) detected per biological replicate for each hair cycle stage examined by ATAC-seq. Nucleosome-free, mono- and di-nucleosome peaks were identifiable for all replicates analysed. **b,** Principal component analysis on reads mapped to peaks separates ATAC-seq biological replicates by hair follicle (HF) cycle stage; *n*=2 (anagen) and *n*= 6 (catagen) **c,** Pearson correlation (R^2^) values for reads mapped to peaks across replicates. **d,** Bar graph of peaks partitioning across the genome. **e,** Heatmap showing all peaks with significantly altered accessibility between anagen and catagen replicates. Accessibility signal for pooled replicate samples is normalized for fraction of reads in peaks, and scaled to reads per genome coverage (RPGC) (colourbar legend, right). Each row corresponds to a detected peak, centered and extended +/- 1kb. Peaks are clustered based on differential accessibility, and summarized as the mean accessibility signal per region in a blue line (peaks gaining accessibility in catagen) or a pink line (peaks losing accessibility in catagen) in the graph at top. **f,** Motif enrichment analysis of the 13,664 peaks losing accessibility in catagen replicates suggests chromatin containing POU-domain transcription factor motifs, among others, is specifically closed relative to the end of anagen. See Supplemental Table 3 for full list.

**Extended Data Fig. 6.**
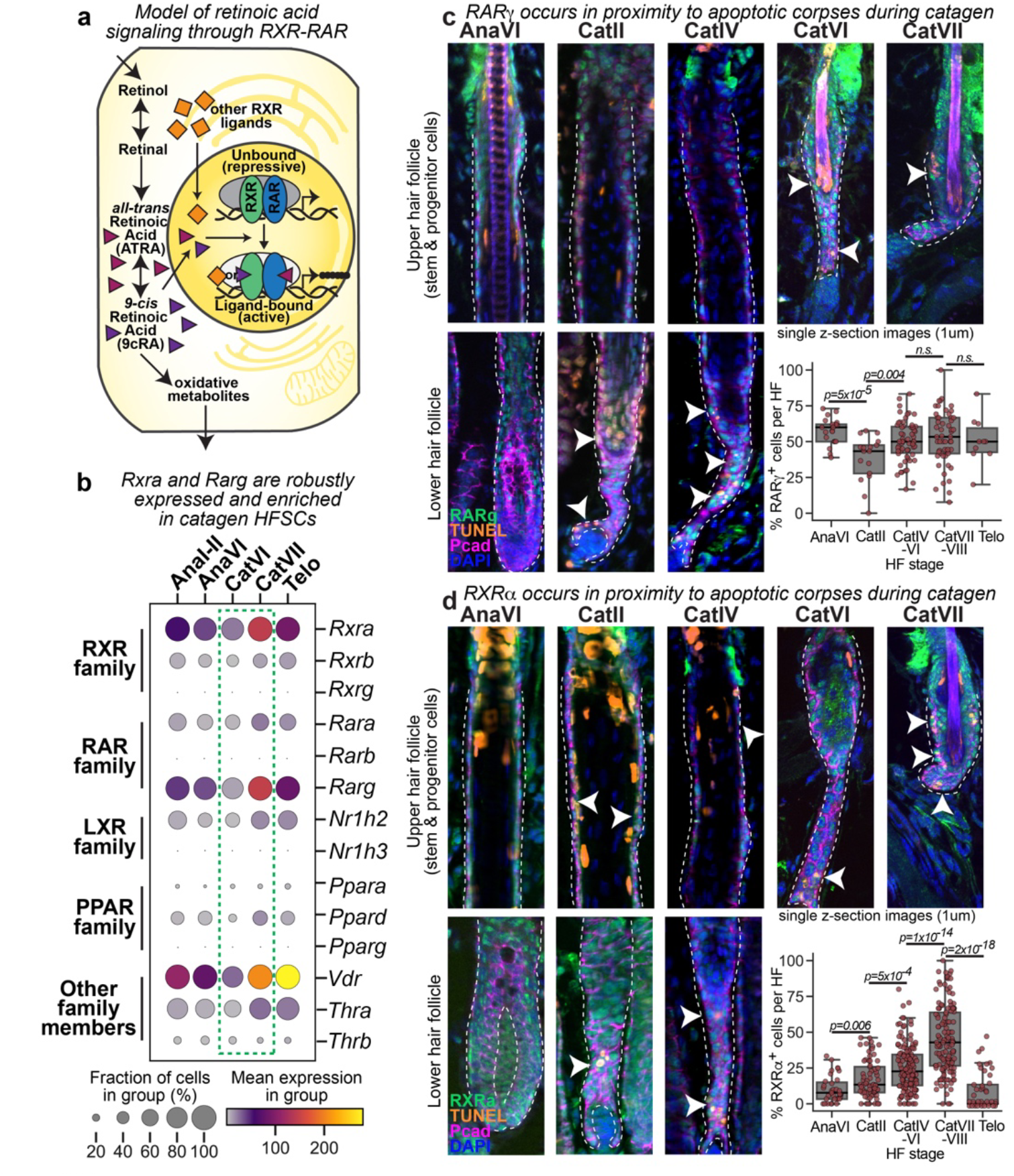
*(Related to Fig. 3)* RXR and RAR family members are expressed in proximity to corpses during apoptotic regression. **a,** Schematic depicting retinoid X receptor (RXR) and retinoic acid receptor (RAR) transcription factor signaling. **b,** Dot plot summarizing non-steroidal nuclear hormone receptor family expression across the hair cycle from single cell transcriptomic data. Each column shows aggregate expression for cells within a hair cycle stage, with dot size corresponding to the percentage of cells within the group expressing the gene and dot colour corresponding to mean normalized counts. **c, d,** Sagittal skin sections of RARγ^+^ cells (**c**) and RXRα^+^ cells (**d**) within hair follicles (dashed white outline) from the end of anagen growth phase (AnaVI) throughout catagen (Cat). Dashed line denotes epithelial-dermal border. Pcad, p-cadherin. Images are representative of 10-20 follicles analyzed per mouse, n=3-6 mice per hair cycle stage. Quantifications based on images above. All quantifications, multiple pairwise Student’s *T*-Tests, p-values indicated. n.s. not significant (p>0.05).

**Extended Data Fig. 7.**
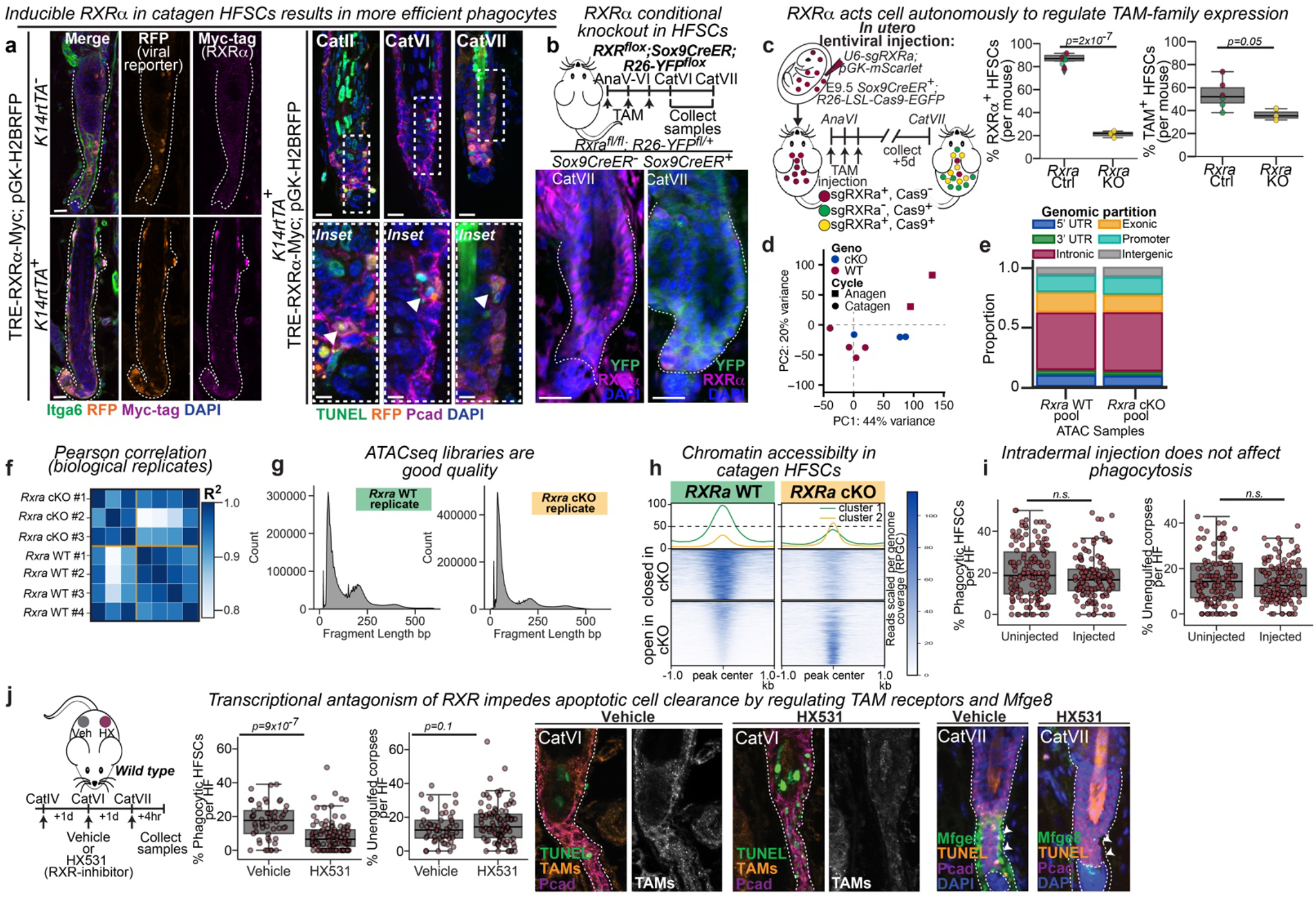
*(Related to Fig. 3)* RXRα regulates the apoptotic cell clearance program *in vivo*. **a,** Sagittal skin sections of *K14rtTA* mice injected with doxycycline (DOX)-inducible *TRE-Rxra-Myc*-tagged lentivirus containing constitutive H2B-RFP infection reporter (TRE-RXRa-Myc;pGK-H2BRFP). Left, DOX treatment induces Myc-tagged RXRα expression only in *K14rtTA*^+^ animals. Scale bar 20um. Right, *K14rtTA*^+^ RFP^+^ (RXRα-high) cells more often contain TUNEL^+^ corpses than non-transduced (RFP^-^) neighbouring cells. Scale bar 20um. Insets, magnified regions of the white dashed box region. Scale bar 10um. Images representative of 10-20 follicles per mouse, *n*=3 mice per genotype per catagen stage. **b,** Top, Experimental strategy. Bottom, Sagittal skin sections of *Rxra* wild type (WT, *Sox9CreER^-^)* and conditional knockout (cKO, *Sox9CreER^+^*) show loss of RXRα in YFP^+^ cKO. **c,** Left, Strategy to mosaically knockout (KO) RXRα by lentiviral injection of single guides against *Rxra* (sgRXRα) with mScarlet transduction reporter into *Sox9CreER^+^; R26-floxed-Cas9-EGFP* mice. Tamoxifen, TAM. Right, Percentages of RXRα^+^ HFSCs (Top) and TAM^+^ HFSCs (Bottom) per mouse. n=3 mice. **d,** Principal component analysis on reads mapped to peaks. **e,** Bar graph of peaks partitioning across the genome. **f,** Pearson correlation (R^2^) values for *Rxra* WT and cKO ATAC-seq replicates. **g,** Representative histograms of fragment lengths in base pairs (bp) for *Rxra* WT and cKO HFSCs. **h,** Heatmap of differentially accessible peaks in late catagen HFSCs between *Rxra* WT and cKO pooled replicates, annotated as in Extended data Fig. 5e. Reads per genome coverage, RPGC; colourbar legend, right. Differential peaks are summarized as mean accessibility signal per region in a yellow line (peaks opening in cKO) or a green line (peaks closing in cKO) in the graph at top. **i,** Percentages of phagocytic HFSCs (left) and unengulfed corpses (right) per late catagen HF (CatVII), either uninjected or intradermally injected with vehicle. n=125-150 HFs across 4-6 mice per condition. **j,** Left, Strategy to transiently inhibit RXR-family transcriptional activity by HX531 intradermal injection. Contralateral backskin was injected with vehicle (Veh) control. Middle, Percentages of corpse-containing HFSCs (Left) and unengulfed corpses (Right) per HF. n=10-15 HFs per mouse; n=8 mice. Right, Sagittal sections of contralateral vehicle- and HX531-injected back skin stained as indicated. Images are representative of n= 10-12 HFs per mouse, n=4 mice per staining panel. All quantifications, multiple pairwise Student’s *T*-Tests, p-values indicated. n.s. not significant (p>0.1).

**Extended Data Fig. 8.**
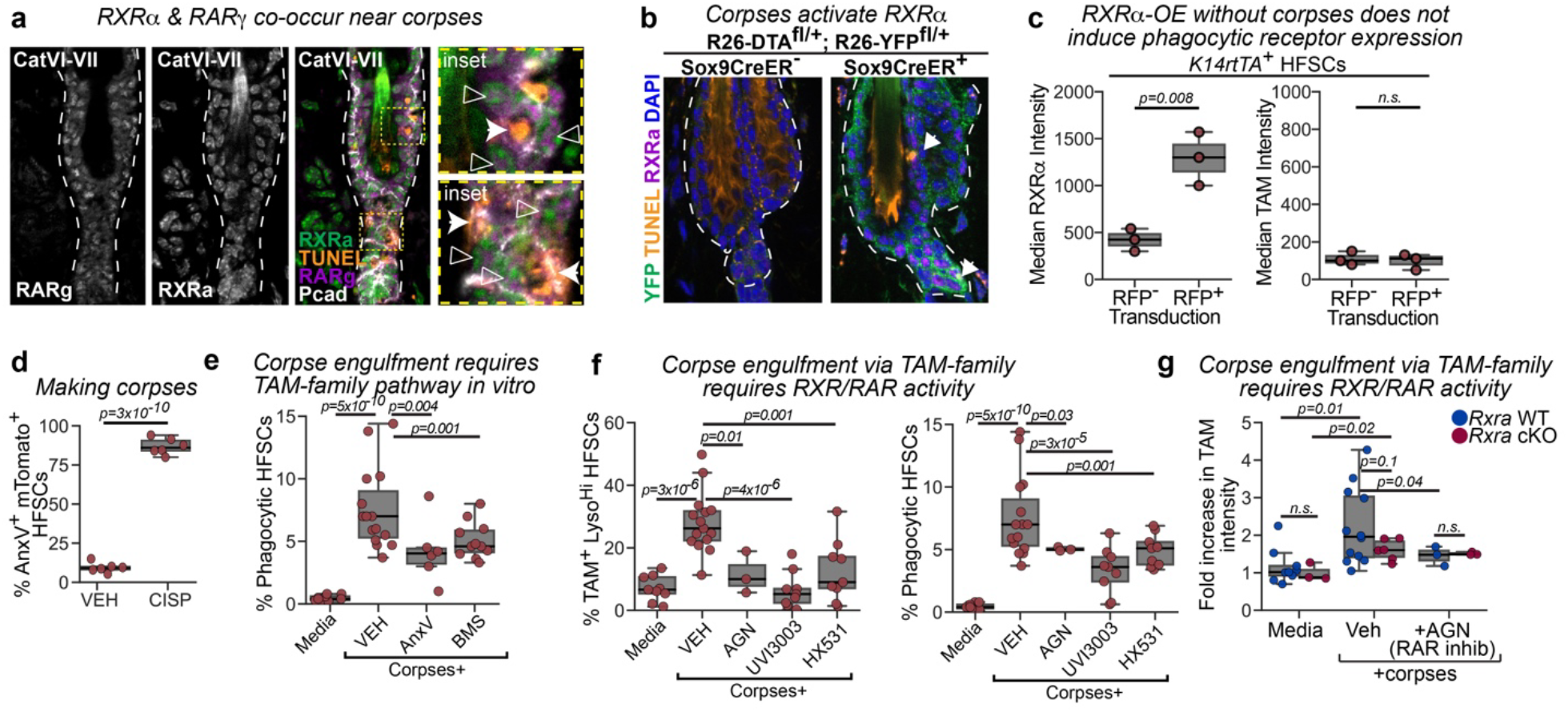
*(Related to Fig. 4)* RXRα responds to corpses. **a,** Representative immunofluorescence images for data in Fig. 4a. Nuclear RXRα and RARγ co-occur (open arrowheads) near unengulfed dying cells (closed arrowheads) in mid-late catagen hair follicles (HFs) (white dashed outline). Insets, higher magnification view of yellow boxed regions to left. 10-20 HFs per mouse, *n*=3-4 animals per hair cycle stage. **b,** Representative immunofluorescence images for FACS-quantified data in Fig. 4b. Diptheria toxin A (DTA) expression in *Sox9CreER^+^* mice creates TUNEL^+^ corpses (white arrowhead) in quiescent HFs (white dashed outline), which induces nuclear RXRα. 10-20 HFs analyzed per mouse, *n*=3-4 animals per genotype. **c,** Quantifications of RXRα intensity (Left) and TAM-family expression (Right) for FACS-isolated hair follicle stem cells (HFSCs) from *K14rtTA*^+^ mice induced to overexpress RXRα (RFP^+^) at the end of catagen and followed to early anagen (AnaII-III), using the same experimental strategy shown in Fig. 3b. *n*=3 mice. **d,** FACS-based percentage of mTomato^+^ HFSCs undergoing apoptosis (AnnexinV^+^) in preparation of corpses/corpse-conditioned media. CISP, cisplatin. *n*=6 replicates. **e,** Percentage of corpse-containing HFSCs, 4 hours post corpse addition, requires phosphatidylserine exposure (blocked by addition of AnnexinV [AnxV]) and TAM-family activity (inhibited by addition of BMS-777607 [BMS]). *n*=6 replicates (AnxV) and n=12 replicates (BMS). **f,** FACS-based quantifications of the percentage of total HFSCs which are TAM-family^+^, lysosome^high^ (left) or contain corpses (right) 4 hours post corpse addition. Inhibition of RAR-family activity by AGN193109 (+AGN), or of RXR-family activity by UVI3003 or HX531. Note, both Media and Corpses+ VEH (vehicle) conditions are the same data as presented in (e), as experiments were carried out together. **g,** FACS-based quantification of TAM-family expression levels, normalized to median expression in media, for *Rxra* wild type (WT) or conditional knockout (cKO) HFSCs 4 hours after corpse addition. All quantifications, multiple pairwise Student’s *T*-Tests, p-values indicated. n.s. not significant (p>0.1).

**Extended Data Fig. 9.**
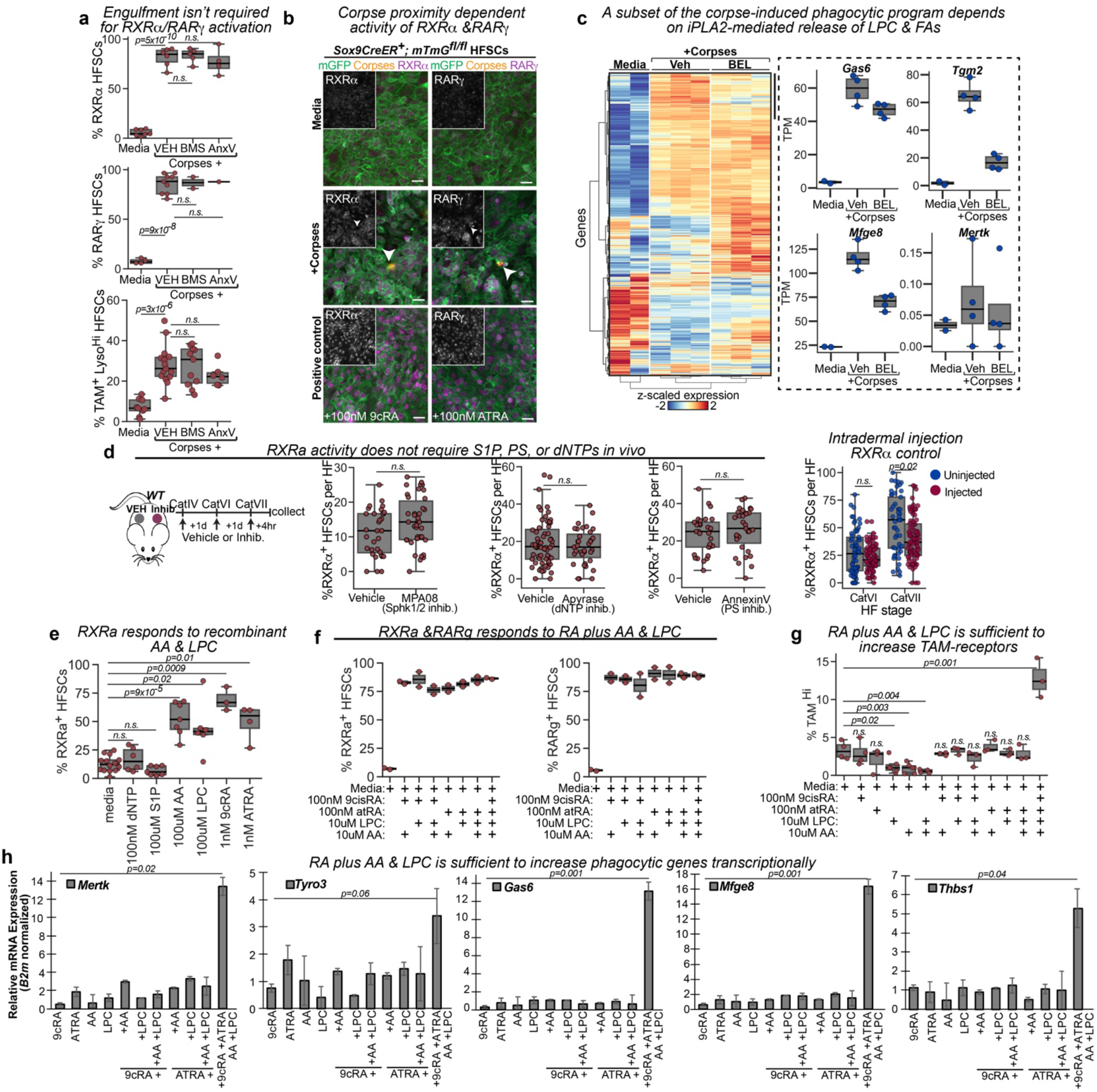
*(Related to Fig. 5)* Corpses secrete lysophosphatidylcholine and free fatty acids to activate RXRα. **a,** Quantifications of RXRα^+^ (Top) and RARγ^+^ (Middle) HFSCs 30 minutes following addition of corpses with (+VEH) or without exposed phosphatidylserine (PS) (+AnxV), or in the presence of TAM-family inhibitor BMS-777607 (+BMS). Bottom, Percentage of TAM-family^+^;Lysosome^high^ HFSCs 4 hours after corpse exposure. *n*=4-8 replicates. **b,** Immunofluorescence of HFSCs stained for RXRα or RARγ 30 minutes after low-titre corpse addition. Insets are RXRα or RARγ alone. Scale bar, 10um. Representative of triplicate experiments performed *n*=3 times. **c,** Left, Heatmap of bulk RNA-sequencing of wild type HFSC replicates exposed to corpses with (+Veh, vehicle) or without (+BEL, bromoenol lactone) secreted lysophosphatidylcholine (LPC) and fatty acids (FAs). Colour bar, Z-score normalized expression. Right, Transcript per million (TPM) expression values of selected phagocytic program genes. Full list in Supplemental Table 5. **d,** Strategy (Left) and percentages of RXRα^+^ HFSCs (Middle) to manipulate corpse-derived sphingosine-1-phosphate (S1P) (left), secreted nucleotides (middle), or exposed phsophatidyleserine, PS, (right) by intradermal injections. Contralateral back skin was injected with vehicle (Veh) control. Right, Uninjected versus vehicle-injected percentage of RXRα^+^ cells. *n*=5-10 HFs per mouse, 3-6 mice per experiment. **e,** Percentage of RXRα^+^ HFSCs in response to recombinant molecules. dATP & dUTP, dNTP; arachidonic acid, AA; 9cRA, 9-*cis* retinoic acid; ATRA, all-*trans* retinoic acid. *n*=4-12 replicates per condition. **f,** Percentage of HFSCs responding to recombinant molecules by increasing RXRα^+^ cells (left) or RARγ^+^ cells (right). *n*=2 replicates per condition, averaged across technical triplicates. **g,** Percentage of TAM-family^+^ HFSCs in response to recombinant molecule combinations. *n*=3-6 replicates per condition. **h,** Quantitative RT-PCR for phagocytic gene transcripts, relative to B2-microglobulin (*B2m*) levels, and normalized to media-only conditions. Concentrations of indicated added molecules are as in (f,g). *n*=3 experimental replicates, performed in technical triplicates. All quantifications, multiple pairwise Student’s *T*-Tests, p-values indicated. n.s. not significant (p>0.1).

**Extended Data Fig. 10.**
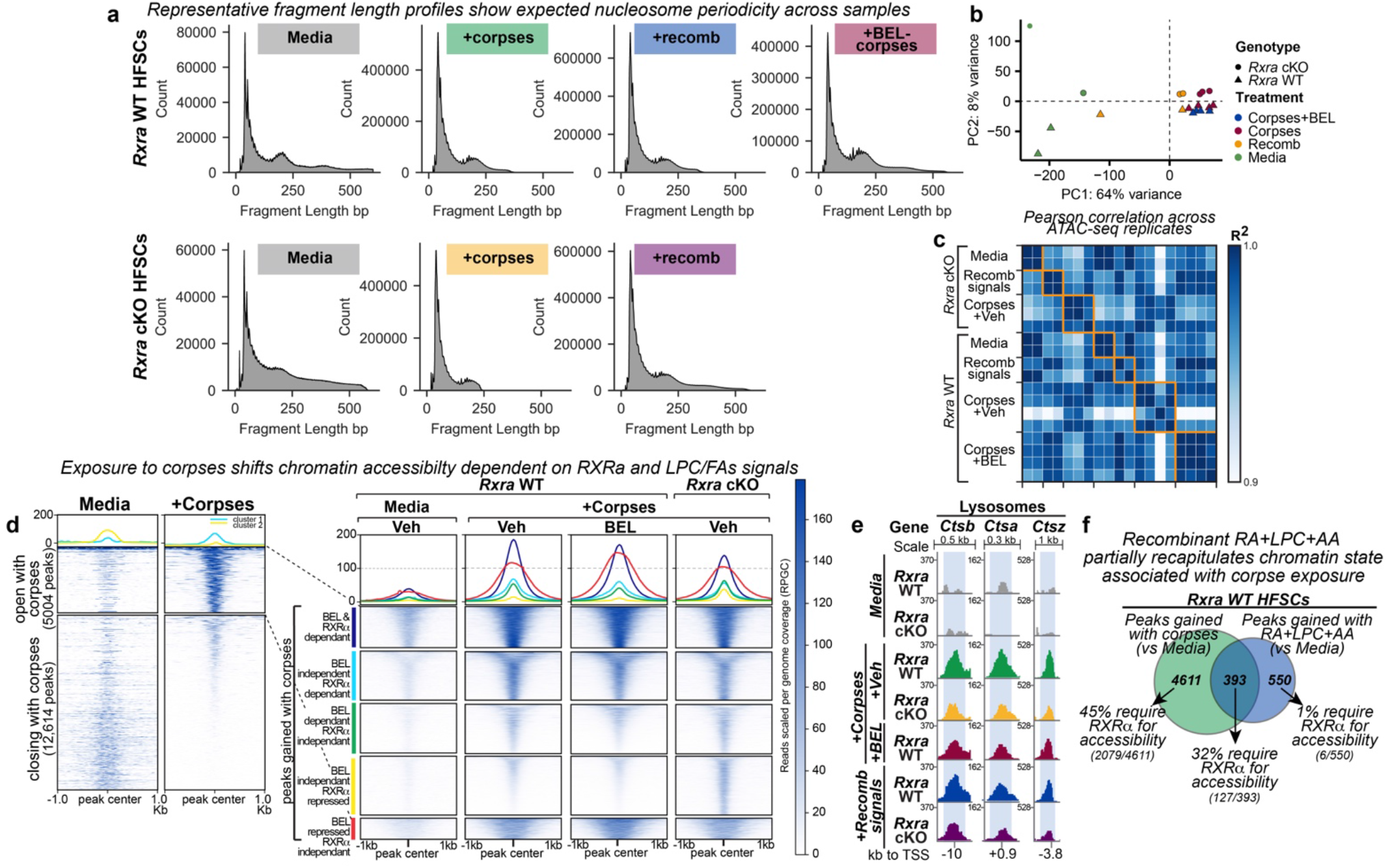
*(Related to Fig. 5)* ATAC-seq shows differential accessibility in response to corpse-derived lysophosphatidylcholine and arachidonic acid dependent on RXRα. **a,** Representative histograms of fragment lengths in base pairs (bp) detected for ATAC-seq replicates. recomb, 9-*cis* retinoic acid + all-*trans* retinoic acid + lysophosphatidylcholine + arachidonic acid; WT, wild type; cKO, conditional knockout. **b,** Principal component analysis on reads mapped to peaks. n=2 biological replicates per genotype per condition. **c,** Pearson correlation (R^2^) values for reads mapped to peaks. **d,** Heatmap showing all peaks with significantly altered accessibility between *Rxra* WT and cKO replicates exposed to corpses with (+Veh, vehicle) or without (+BEL, bromoenol lactone) lysophosphatidylcholine + arachidonic acid. Signal for pooled replicate samples is in reads per genome coverage (RPGC) (colour bar legend, right). Each row corresponds to a detected peak, centered and extended +/- 1kb. Peaks are clustered based on differential accessibility relative to *Rxra* WT+Veh corpses, and summarized as the mean accessibility signal per region in a dark blue line (peaks that close in BEL and *Rxra* cKO), a light blue line (peaks that close in cKO only), a green line (peaks that close in BEL only), a yellow line (peaks that open in cKO only) or a red line (peaks that open in BEL only) in the graph at top. **e,** Replicate pooled peak tracks for enhancers associated with lysosomal genes. Annotated as in Fig. 5f. **f,** Venn diagram representation comparing peaks that open in response to corpses with those gained in response to retinoic acid (RA) + lysophosphatidylcholine (LPC) + arachidonic acid (AA) (‘Recomb’ condition) in *Rxra* WT HFSCs. Arrows indicate the percentage of peaks that require RXRα for accessibility within each category. Notably, enhancer peaks associated to phagocytic genes fall within the 32% of peaks gained in response to both corpses and recombinant signal that require *Rxra* for accessibility, which are shown in Fig. 5f.

## Methods

The following previously generated mouse lines were used in this study: *Rxra^Flox^* (ref.^55^; Jax stock 013086)*, Sox9CreER* (ref.^56^)*, Krt14-rtTA* (ref.^57^; Jax stock 008099)*, Rosa26^lox-STOP-lox-YFP^* (ref.^58^; Jax stock 006148; referred to as *R26^YFP^*)*, Rosa26^mTmG^* (ref.^59^; Jax stock 007576; referred to as *R26^mTmG^*)*, Rosa26^Brainbow2.1^* (ref. ^60^; Jax stock 013731, referred to as *R26^Brainbow2.1^*)*, Rosa26^lox-STOP-lox-Cas9-EGFP^* (ref.^61^; Jax stock 026175, referred to as *R26^Cas9-EGFP^*)*, Rosa26^lox-STOP-lox-DTA^*(ref.^62^; Jax stock 010527, referred to as *R26^DTA^*) and *Mertk^-/-^* (full knockout, ref.^63^). The *Mertk* knockout mice used in this study are referred to as *Mertk^-/-V2^* in the originating paper. Wild type CD1 or C57BL/6 animals were originally purchased from Charles River and The Jackson Laboratories, respectively, and maintained as in house colonies.

Mice were maintained and bred under specific-pathogen-free conditions at the Comparative Bioscience Center (CBC) at The Rockefeller University, an Association for Assessment and Accreditation of Laboratory Animal Care (AALAC)-accredited facility. *Mertk* knockout animals and C57BL/6J wild type controls (maintained as separate colonies) were bred and maintained in a specific-pathogen-free facility at Yale University. All animals were bred and maintained under a strict 12-hour light cycle and fed with standard chow. The temperature of the animal rooms was 20-26°C, and the humidity was 30-70%. Adult animals were housed in cage with a maximum of five mice. All mouse protocols were approved by the Institutional Animal Care and Use Committee (IACUC) at The Rockefeller University, or by the IACUC at Yale University.

For comparative assessments of phenotype between control and mutant animals, age and sex matched mice were used, with preference given to littermate controls wherever possible. For our inducible overexpression studies, a *Krt14-rtTA*^+/-^ (heterozygous) male was mated with CD1 females and all offspring were transduced with lentivirus at embryonic day (E) 9.5 (see following sections). Offspring of both genotypes received doxycycline by intraperitoneal (i.p.) injection (0.5mg/mouse) at postnatal day (P) 14 to activate *Krt14-rtTA* within 12hr, and expression was maintained by feeding the mother and pups doxycycline (2mg/kg) chow (Bioserv). *Krt14-rtTA*^-^ animals were used as control, with *Krt14rtTA*^+^ mice as the experimental group. To generate RXRa control and conditional knockout mice for experiments, the *RXRa^Flox^* line was crossed with *Sox9CreER^+^; R26^YFP^* animals*. Sox9CreER*^-^ mice with any *Rxra^Flox^;R26^YFP^*genotype, and *Rxra^fl/+^;Sox9CreER^+^;R26-YFP^fl/+^*mice were used as controls, while experimental mice were *Rxra^fl/fl^;Sox9CreER^+^;R26-YFP^fl/+^.* All animals received tamoxifen (2% in corn oil) (Sigma-Aldrich) to activate *Sox9CreER,* administered by i.p. injection once a day for 3 days, as indicated. *Sox9CreER* was similarly activated when crossed to *R26^mTmG^* (to label HFSCs prior to FACS-isolation and culture)*, R26^Brainbow2.1^* (to stochastically label HFSCs and identify functional phagocytes), and *R26^Cas9-EGFP^*(to mosaically knockout *Rxra*). To activate *Sox9CreER* sparsely when crossed to *R26^DTA^*, 2% tamoxifen was i.p. injected once early in second telogen.

### Hair cycle staging

Male and female mice have different hair cycle lengths due to a longer telogen quiescence phase in females, but otherwise progress through the hair cycle similarly. In addition to sex, strain and individuals also affect hair cycle stages. Therefore, we always determine hair cycle stage by visual inspection, and morphological staging on sectioned tissue. Specifically, for C57BL/6 pure and mixed backgrounds, visual inspection was performed by trimming full-length telogen hairs with electric clippers to reveal dorsal skin. Hair follicle (HF) entry into anagen was determined by darkening of skin and reappearance of hair. Catagen progression was determined by lightening of the skin, which appears black at the end of anagen, to a near complete loss of pigmentation (greyish-pink skin) by late catagen. Entry into telogen was marked by the appearance of completely unpigmented (pink) skin. In unpigmented mice (CD1 strains), histological analysis of HF morphology was used to confirm hair cycle staging based on relative age.

For all experiments, a small piece of midline dorsal skin was taken in parallel from each mouse, fixed and processed for sectioning and immunofluorescence to precisely stage the hair cycle. Hair cycle was staged based HF morphology^64-67^, as well as by immunofluorescence for markers of anagen (EdU-incorporation following a 2hr pulse chase), catagen (cleaved caspase-3 and/or TUNEL positivity) or telogen (pSMAD1/5/9 and/or Lef1). For samples obtained from anagen or catagen stage mice the dorsal back skin was subdivided in two along the anterior-posterior axis prior to the experiment, as precise hair cycle stage differs anterior to posterior (with anterior generally one substage ahead). For anagen samples, 5’-ethynyl-2’-deoxyuridine (EdU) was injected i.p. (50ug/g) (Sigma-Aldrich) and chased for 2hr prior to collection.

### Induction and knockout constructs

#### RXRα-induction

To make TRE-RXRa-Myc; pGK-H2B-RFP, human *RXRa* cDNA was PCR-amplified from pSV-Sport-RXRα (a gift from Bruce Spiegelman; Addgene #8882)^68^, and a NheI site was introduced at the 5’end. This was then inserted into the NheI and EcoRI restriction sites of a pLKO vector modified to contain the inducible tetracycline response element at the 5’ end, as well as a 3’-myc epitope tag. Prior to packaging this construct as a lentivirus, induction of *RXRa* was tested in culture using FACS-isolated *Krt14rtTA*^+^ keratinocytes grown in E300 media. Briefly, cells were transiently transfected with the TRE-RXRα construct using Effectene into keratinocytes in a 6-well plate format, following the manufacturer’s protocol (Invitrogen). 48 hrs later doxycycline (100ng/ml) was added to induce RXRα expression for a further 24 hrs. Cells were fixed and stained for Myc-tag and RXRa, as described for cell culture immunofluorescence.

#### RXRα-mosaic knockout

To identify efficient CRISPR guide RNAs (sgRNA) against mouse *Rxra*, we synthesized oligos targeting exon 4 with BsmBI restriction sites at 5’ and 3’ respectively (IDT). Oligos were subcloned into pLentiGuide-Puro (a gift from Feng Zhang; Addgene #52963), following the Zhang lab protocol^69^. To select a guide for *in vivo* use, we first tested the cutting efficiency in culture using *K14Cre*^+^; *R26^Cas9-EGFP^* expressing keratinocytes. pLentiGuide-Puro constructs were transiently transfected using Effectene into keratinocytes in a 6-well plate format, following the manufacturer’s protocol (Invitrogen). After 72 hrs, genomic DNA was harvested using QuickExtract DNA Extraction Solution (Lucigen), and gDNA was prepared by heating to 65°C for 10 min followed by heat inactivation at 95°C for 2 min. Following PCR-amplification of each guide target region, a T7 endonuclease I cutting assay (NEB) was used to identify the extent of insertions/deletions for each guide. The most efficient guide showed ∼70% genome editing events *in vitro*, and was cloned with its U6 promoter into a modified pLKO vector containing a constitutive pGK-driven mScarlet fluorophore (3’ to the sgRNA) for lentiviral preparation.

### Lentiviral preparation and Injection

High-titre lentivirus was prepared and embryonic day (E) 9.5 embryos of indicated genotypes were infected with lentivirus delivered by ultrasound-guidance microinjection into the amniotic sac as previously described^70,71^. At E9.5 the surface ectoderm exists as a single layer of unspecified skin progenitors, which can be efficiently, selectively and stably transduced by the viral DNA, without transduction of dermal cell types^71^.

### HFSC culture

All primary hair follicle stem cell (HFSC) lines were grown on a layer of mitomycin C-inactivated 3T3/J2 feeder fibroblast cells, and maintained in E intermediate (300uM) calcium media^72^ supplemented with 10uM Y-27632 (Selleckchem) (“E300-Y media”)^73^. The 3T3/J2 fibroblast cell line was expanded in DMEM/F12 media (Thermo Fisher Scientific) with 10% CFS (Gibco), 100U/ml streptomycin and 100mg/ml penicillin. Cells were grown at 37°C, with 7.5% CO2, and media was routinely changed every 2-3 days. Cell lines were grown to confluency, then propagated by digesting with 0.25% Trypsin EDTA (Gibco) for 5-10 min at 37°C and resuspended with culture media for passaging. Experiments were conducted with cells at passages 8-10. For experiments, cells were switched to E intermediate calcium media without Y-27632 (“E300 media”) and cultured for 24-48hr prior to the experiment.

Primary HFSCs were derived from the following mouse crosses at second telogen: *R26-mTmG^fl/+^; Sox9CreER^-^* (mTomato^+^ HFSCs to make apoptotic corpses), *R26-mTmG^fl/+^; Sox9CreER^+^*(mGFP^+^ HFSCs to make naïve HFSC to expose to corpses), *Rxra^+/+^; Sox9CreER^+^; R26-YFP^fl/fl^* (*Rxra* wild type YFP^+^ HFSCs), and *Rxra^fl/fl^; Sox9CreER^+^; R26-YFP^fl/fl^* (*Rxra* conditional knockout YFP^+^ HFSCs). All HFSCs were FACS-isolated (described later) and cultured as described. To generate *Rxra* wild type and conditional knockout HFSC lines, cells were FACS isolated, and cultures were established prior to activation of *Sox9CreER by* 4- hydroxytamoxifen (4-OHT). At passage 2, *Sox9CreER* was activated in culture by 4-OHT in solution (Sigma Aldrich); to do so it used at a final concentration of 1uM in E300-Y media to treat HFSCs. Media plus 4-OHT was refreshed each day for three consecutive days, and then replaced by E300-Y. HFSCs were allowed to grow for 4 further days prior to FACS isolation of YFP^+^ cells.

To prepare corpses, fully confluent mTomato^+^ HFSCs were treated with 200uM cisplatin (in 0.9% saline) for 18hrs. Dead and dying cells were collected from the supernatant by pelleting at 700xg for 5 min, washed once with E300 media, and returned to the plate in fresh E300 media. Prior to returning the floating corpses, dying adherent cells were rinsed with PBS to remove residual cisplatin and a minimal amount of fresh E300 media was added. Corpses were allowed to condition the media for a further 3-4hrs. Apoptotic cell corpses were collected by tapping the side of the plate and pipetting their media over them to detach dying cells. Floating and detached corpses were collected and pelleted by centrifugation as before. Corpse conditioned media was carefully removed to a separate tube, before resuspending corpses in a minimal volume of fresh E300 media. To label any corpses derived from 3T3/J2 fibroblasts, corpses were next incubated with DiI-CM (Invitrogen) for 5 min at 37°C before pelleting and washing with PBS as before. DiI/mTomato^+^ corpses were resuspended in their corpse conditioned media, counted and aliquoted (in their conditioned media) directly onto experimental plates (media removed prior) at a ratio of roughly 10 corpses:1 HFSC. For corpse-conditioned media experiments, corpses were prepared as described and then spun out of the media at 1200xg for 10 min. Corpse-conditioned media was further strained through a 0.45um syringe filter, prior to use. Corpses and/or conditioned media were always prepared immediately prior to their use.

Manipulation of corpse-derived signals was achieved by adding small molecule inhibitors to the corpses after the removal of cisplatin (16uM bromoenol lactone (Sigma Aldrich); 100nM MPA08) or by incubating the corpses with recombinant molecules for 15-20 min prior to adding them to naïve HFSCs (1U Apyrase; 1ng/ml AnnexinV). Vehicle treated control corpses were incubated with 1% DMSO. Similar preparations were made for corpse-conditioned media. To manipulate corpse-sensing mechanisms on HFSCs, naïve mGFP^+^ or YFP^+^ HFSCs were pretreated with the indicated antagonist (1nM UVI3003; 1uM HX531; 1uM JTE013; 100nM BMS 777607; 1uM AGN 193109) (first four: Tocris Bioscience; last one: R&D Systems) for 30 min prior to corpse or corpse-conditioned media addition. When adding corpses or conditioned media, the concentration of antagonist was maintained by adding an additional amount of the appropriate compound to the corpses and/or conditioned media.

To test the ability of recombinant molecules to recapitulate corpse secreted signals, mGFP+ or YFP+ HFSCs were cultured in E300 media plus the indicated concentrations of recombinant molecules (see figures and figure legends). Molecules were prepared and stored as stock solutions according to manufacturer’s instructions. Briefly, 9-*cis* retinoic acid and all-t*rans* retinoic acid (both R&D Systems) were each dissolved in 100% DMSO, protected from light, and stored long term at -80°C with working solutions kept at -20°C. Recombinant lysophosphatidylcholine, sphingosine-1-phosphate, and arachidonic acid (all Tocris Bioscience) were each dissolved in 100% ethanol and stored like the retinoids. Free nucleotides were purchased as 100mM stocks of dATP or dUTP as sodium salts in ultrapure water (NEB) and stored at -20°C. Stock solutions were diluted individually or in combinations with E300 media for experiments.

In pilot experiments nuclear accumulation (by immunofluorescence) of RXRα or RARγ peaked at 30 min post corpse exposure, and so that time point was used to assess immediate effects of corpse-derived signals or recombinant molecules in subsequent experiments. Similarly, the number of corpse-containing HFSCs plateaued at 4-6 hrs after corpse addition, and so transcriptional activation of the phagocytic program, surface expression of phagocytic receptors and corpse engulfment (latter two by FACS) were routinely assessed at that time point.

### Intradermal injections

Adult mice in early second catagen (CatII) were anesthetized using isoflurane prior to intradermal injections. Isofluorane anesthetization was maintained throughout the procedure using a nose cone for delivery. Back skin was shaved using electric clippers and the surface sterilized by wiping with ethanol wipes. Vehicle or small molecule containing solutions were prepared by diluting appropriate chemical in sterile PBS plus 1% FluoSpheres Carboxylate-modified microspheres (1.0um, Thermo Fisher Scientific #F8816) to assess placement of the injection on tissue sections. To find the injection site for repeated intradermal injections on subsequent days, a small dot was made with permanent marker which the needle was inserted through. 1ml insulin syringes with the needle bent to approximately 45° were used to shallowly inject through the epidermis to the dermal space approximately 3-5mm from the injection site. An injection volume of 25ul was delivered per injection site, with an average of 4 injection sites per mouse: two anterior and two posterior. Vehicle injections (10% DMSO) were randomly designated to either the left or right side, with the contralateral skin receiving the indicated small molecules. Following the injections, mice were placed in their home cage on a heating pad to recover. Compounds were prepared at 100X the cell culture working solution, from the same stock solutions. To inhibit the phagocytic program across the course of catagen, intradermal injections were performed three times, separated by 20-24hrs. Animals were sacrificed by lethal CO2 administration, 4hrs after the final injection (at CatVII-VIII).

### Tissue collection and sectioning

For immunofluorescence analysis of tissue sections, mice were shaved following lethal CO2 administration and their back skin dissected. Back skin was stretched onto Whatman paper for stability, and immediately prefixed in 1% or 4% paraformaldehyde (PFA) for 1 hr at 4°C or 30 min at 25°C, respectively. After fixing, tissue was washed twice with PBS for 10 min at 4°C, before incubating in 30% sucrose in PBS at 4°C overnight. Tissue was embedded in OCT medium (VWR) and frozen on dry ice blocks before storage at -80°C. Alternatively, fresh frozen tissue was prepared without prefixation by directly embedding the skin in OCT after it was placed on whatman paper. Frozen tissue blocks were sectioned at 20um on a Leica cryostat and mounted on SuperFrost Plus slides (Thermo Fisher). When necessary, sections were stored at -20°C prior to use.

### Immunofluorescence

#### Skin sections

Following sectioning, tissue was allowed to dry on the slide for 1 hr in a partially closed slide box. Fresh frozen tissue was post-fixed with 4% PFA for 5 min, followed by washing in phosphate- buffered saline (PBS) three times for 5 min each. Pre-fixed tissue sections started with the PBS wash step to remove attached Whatman paper. Following washes, samples were permeabilized and blocked in blocking buffer (5% donkey serum, 2.5% fish gelatin, 1% BSA, 0.3% Triton in PBS) for 1 hr at room temperature. Primary antibodies were incubated overnight at 4°C, samples were washed for 5 min in PBS (three times) at room temperature, and secondary antibodies were incubated together with DAPI (to label nuclei) for 1 hr at room temperature. Following three final PBS washes of 5 min each, samples were mounted in Prolong Diamond Antifade Mountant (Invitrogen) for imaging. For TUNEL labelling, the Cell Death Detection Kit (TMR red or FITC; Roche) was used according to manufacturer’s instructions, with application of secondary antibodies. A modification was made to halve the concentration of the substrate labelling component to reduce background fluorescence in the skin. Antibodies were used as follows: rabbit anti-cleaved-caspase-3 (Cell Signalling, 9661, 1:250), rat anti-RFP (Chromotek, 5F8, 1:1000), rabbit anti-RFP (MBL, PM005, 1:1000), chicken anti-GFP/YFP (Abcam, ab13970, 1:1000), goat anti-P-cadherin (R&D, AF761, 1:250), rabbit anti-keratin14 (Fuchs laboratory, 1:200), rabbit anti-myc epitope (71D10) (Cell Signalling, 2278, 1:250), rat biotinylated anti-CD45 (Biolegend, 5530, 1:200), rabbit anti-RXRα (D6H10) (Cell Signalling, 3085, 1:250), rabbit anti-RARγ (D3A4) (Cell Signaling, 8965, 1:250), rabbit anti-MFGE8 (Invitrogen, PA5-109955, 1:200), rat AlexaFluor647-conjugated anti-F4/80 (BM8) (Biolegend, 123121, 1:200), and rat biotinylated anti-Itga6/CD49f (GoH3) (Biolegend, 313603, 1:500). All secondary antibodies used were raised in a donkey host, and conjugated to AlexaFluor350, AlexaFluor488, Rhodamine, AlexaFluor568, or AlexaFluor647 (Jackson ImmunoResearch Laboratory; 1:500). 4’,6-diamidino-2- phenylindole (DAPI) was used to label nuclei (1:10,000). To co-stain RARγ and RXRα, the rabbit primary antibodies were individually directly conjugated to one of AlexaFluor350, AlexaFluor488, AlexaFluor568, or AlexaFluor647 using the rabbit specific Zenon Antibody Labelling Kit (Thermo Fisher Scientific) and following manufacturer’s instructions.

#### Cell culture

For immunofluorescence experiments, feeders were split onto poly-L-lysine coated glass coverslips, seeded in 12-well plates 24 hrs prior to the addition of HFSCs. HFSCs were grown to confluency before feeders were detached by repeated PBS washes, and corpse or corpse conditioned media experiments were performed. At the end of the experiment, cells were washed twice with PBS and prefixed with 4% PFA for 3 min at 25°C. Cells were washed three times with PBS and stained as for tissue sections.

### Microscopy

Images of *Sox9CreER; Rosa26^Brainbow2.^*^1^ tissue was acquired using a Zen-software driven Zeiss LSM 780 inverted laser scanning confocal microscope and 20X air objective (NA=0.8), a 40X water immersion objective (NA=1.2), or a 63X oil immersion objective (NA=1.4). To separate CFP, YFP, GFP, RFP and AlexaFluor647 fluorophores, excitation with specific laser lines (405, 440, 488, 514, 561, 594, and 633) and narrow wavelength emission cut-offs on 4 detectors were set up as follows: CFP (ex.440nm, em.450nm- 490nm), GFP (ex. 488nm, em. 500nm-515nm), YFP (ex. 514nm, em. 525nm-570nm), RFP (ex. 561nm, em. 595nm-620nm), and AlexaFluor647 (ex. 633nm, em. 650nm-690nm). Due to their well-separated excitation and emission spectra, GFP and AlexaFluor647 were acquired simultaneously on the same detector. Stacks with a 1um step were acquired. Confocal microscopy was performed in The Rockefeller University’s Bio- Imaging Resource Center, RRID:SCR_017791.

Images of all other cryosections were acquired using a Zen-software driven Zeiss Axio Observer.Z1 epifluorescent/brightfield microscope with a Hamamatsu ORCA-ER camera, Axiocam350, and an ApoTome.2 slider (to reduce light scatter in *z*). Stacks with a 1um step were acquired. Apotome acquired images were processed via “Apotome Raw Convert” function, and stitched (if necessary), in Zen software. Subsequent image processing was conducted in ImageJ. For presentation purposes, images were cropped and assembled in Adobe Illustrator.

### Electron microscopy

Dissected back skin was placed on thin paper towel for stability, and fixed in 2% glutaraldehyde, 4% PFA, and 2mM CaCl_2_ in 0.1 M sodium cacodylate buffer (pH 7.2) for 2hr at room temperature, postfixed in 1% osmium tetraoxide and processed for Epon embedding. Ultrathin sections of 60-65nm were counterstained with uranyl acetate and lead citrate, before images were taken with a transmission electron microscope (Tecnai G2-12;FEI) equipped with a digital camera (AMT BioSprint29). Samples were processed and imaged at The Rockefeller Electron Microscopy Resource Center.

### Flow cytometry

To obtain single-cell suspensions for fluorescence activated cell sorting (FACS) at all stages of the hair cycle, back skin was excised, and the dermal side scraped with a dull scalpel to remove excess fat prior to incubation with 0.25% collagenase (Sigma-Aldrich) in warm PBS, dermal side down for 45-60 min at 37°C with gentle rotation in a plastic petri dish. The dermal side was scraped gently with a dull scalpel to mechanically dissociate cells in the lower outer root sheath (ORS) and hair bulb (“dermal fraction”). The dermal fraction was only kept for late anagen and early-mid catagen samples, and was processed separately from the epidermal fraction. To collect the epidermal fraction, the skin was placed dermal side down in 0.25% trypsin-EDTA (Gibco) for 20-25 min at 37°C with gentle rotation. The hairy side of the skin was scraped against the direction of hair growth with a dull scalpel to release cells in the upper HF (including the HF bulge stem and hair germ progenitor cells). For both dermal and epidermal fractions, the resulting cell suspensions were pipetted up and down with a 5ml serological pipette for 5 minutes, before being quenched with FACS buffer (5% fetal bovine serum, FBS, in PBS). Plastic petri dishes were rinsed with 5ml of FACS buffer 2-3 times, which was collected and added to the appropriate cell suspension. Suspensions were filtered through sequential 70um and 40 um nylon filters (VWR), before being pelleted at 350xg for 15 min at 4°C. Cell pellets were resuspended in ice cold FACS buffer, re-filtered into FACS tubes, and incubated with primary antibodies for 20 min on ice. Secondary antibodies and LysoTracker DeepRed (Invitrogen, 1:4000) were added directly to FACS tubes, and incubation continued for 10 min on ice. Samples were further diluted with FACS buffer plus DNase (Roche) to minimize cell clumping prior to sorting or analysis. For analysis of RXRα levels by FACS, cells were stained with cell-surface specific primary and secondary antibodies, before being fixed and processed using the BD Cytofix/Cytoperm kit following manufacturer’s instructions. Primary antibodies were used as follows: rat biotinylated anti-CD45 (30-F11) (eBioscience, Cat #13-0451-82, 1:200), rat biotinylated anti-CD117 (2B8) (eBioscience, 13-1171-82, 1:200), rat biotinylated anti-CD140a (APA5) (eBioscience, 13-1401-82, 1:200), rat biotinylated anti-CD31 (390) (eBioscience, 13-0311-82, 1:200), rat anti CD34-FITC (RAM34) (eBioscience, 11-0341-82, 1:200), rat anti CD34-eFluor660 (RAM34) (eBioscience, 50-0341-82,1:200), rat anti CD49f/Itga6-PercpCy5.5 (GoH3) (BioLegend, 313617, 1:250), rat anti-Ly6A/E(Sca1)-APC-Cy7(BioLegend, 108125, 1:1000), rabbit anti-RXRα (D6H10) (CST, 3085, 1:250), rat anti-Tyro3/Dtk-AlexaFluor700 (R&D Systems, FAB759N, 1:200), rat anti-Mertk-AlexaFluor700 (R&D Systems, FAB5912N, 1:200), and rat anti-Axl-AlexaFluor700 (R&D Systems, FAB8541N, 1:200). Secondary antibodies were used as follows: Strepavidin-PE-Cy7 (1:3000) and donkey AlexaFluor 488 or AlexaFluor568 (1:500). AnnexinV-AlexaFluor568 (Invitrogen, A13202, 1:100) and/or DAPI was used to identify apoptotic and dying cells, respectively. For FACS using AnnexinV, primary and secondary antibody staining was performed in AnnexinV Binding Buffer (10mM HEPES, 140mM NaCl, 2.5mM CaCl_2_, pH 7.4).

For HFSC isolation for *in vitro* culture and bulk ATAC-sequencing, cells were sorted using a 70um nozzle into E300-Y media and FACS buffer. For bulk RNA-sequencing, cells were sorted using a 70um nozzle directly into Trizol and FACS buffer. For single cell RNA-sequencing cells were sorted with an 85um nozzle into 96-well PCR plates (Bio-Rad) containing 2ul of lysis buffer [0.2% Triton X-100, 2U/ul RNaseOUT (Thermo Scientific), 0.25uM oligo-dT30VN primer, 1:2×10^6^ diluted ERCC spike-in RNAs (Ambion)]. Sorting was performed on a BD FACSAriaII equipped with Diva software (BD Biosciences).

For FACS analysis, live cell suspensions from back skin were collected and analyzed in the same manner as for sorting. Alternatively, cultured HFSCs were trypsinized for 7-10min (as for passaging the cell lines), and pelleted at 300xg before resuspension, filtering and incubating with primary antibodies. A minimum of 20,000 HFSCs were analyzed per sample using either a BD LSRII Flow Cytometer or a BD Fortessa Flow Cytometer (BD Bioscience). For analysis of the Brainbow2.1 HFSCs, a LSRII specially equipped with a 445nm laser was used to excite CFP, separately from YFP/GFP and RFP.

Flow cytometry was performed at The Rockefeller University’s Flow Cytometry Resource Center, RRID:SCR_017694. Flow cytometry plots were generated using FlowJo to illustrate the strategies used for cell isolation, and manually compensated for presentation. Representative sort schemes pertaining to *Sox9CreER*; *R26^Brainbow2.1^* analysis of functionally phagocytic HFSCs, wild-type isolation of HFSCs for single- cell RNAseq and ATACseq across the hair cycle, RXRα overexpression and knockout in HFSCs, and *Sox9CreER;R26^DTA^*ectopic corpse response are available in Supplementary Information.

### scRNA-sequencing libraries

Single-cell RNA-sequencing libraries were prepared from FACS-isolated HF epithelial cells in AnaVI, CatVI, and CatVII, using a slightly modified Smart-Seq2 protocol as previously described^74,75^. For each hair cycle stage, cells from 3-6 animals were pooled prior to FACS-isolation. In brief, cells were sorted into hypotonic lysis buffer, snap frozen in liquid nitrogen and stored at -80°C until all samples were collected. Cells were lysed by heating at 72°C for 3 min, followed by reverse transcription of mRNA using dT30 oligos, template switching oligos and Maxima H- reverse transcriptase. The whole transcriptome was amplified (15 cycles) by KAPA HiFi DNA polymerase (Roche), and then size-selected using 0.6X AmpPure XP beads (Beckman Coulter). To exclude cells with poor amplification, and wells containing multiple cells, qPCR for *Gapdh* was performed. Illumina sequencing libraries were indexed with unique 5’ and 3’ barcode combinations (up to 384 cells) using the Nextera XT DNA library preparation kit (Illumina). Libraries were pooled and size-selected with 0.9X AmpPure XP beads. Prior to sequencing on Illumina NextSeq500 using a 75 bp paired-end read mid-output setting, library quality was assessed by TapeStation (Agilent).

### ATAC-sequencing libraries

ATAC-seq was performed on 20,000-75,000 (*in vivo* samples) or 50,000 (culture samples) FACS-sorted HFSCs, as previously described^76-78^. Briefly, cells were lysed in ATAC lysis buffer for 1 min on ice, washed and nuclei resuspended in transposase buffer. Genomic DNA was transposed using Tn5 transposase (Illumina) for 30 min at 37°C, at which point the reaction was halted. Samples were uniquely barcoded in batches of 10-12 (in vivo samples) or one batch of 27 (cultured HFSCs), using Buenrostro^76^ or Nextera XT index kit v2 indices. Sequencing libraries were prepared according to manufacturer’s instructions (Illumina). Libraries were sequenced to a depth of 50-100 million sequences, using paired-end runs on an Illumina Novaseq 6000 (at The Rockefeller University Genomics Resource Center).

### RNA isolation

Total RNA was isolated from FACS-isolated HFSCs using the Direct-zol RNA MicroPrep kit (Zymo Research) following manufacturer’s instructions. The optional DNase I treatment was included in all sample preps, and RNA was eluted in DNase/RNase-free water. Quality and concentration of RNA samples were determined using an Agilent 2100 Bioanalyzer. All samples for sequencing had RNA integrity (RIN) numbers >8.5. RNA samples were used for RT-qPCR or bulk RNA-sequencing, as described.

### Bulk RNA-sequencing libraries

Comparable amounts of RNA per sample were used to prepare bulk RNA-sequencing libraries using Illumina Trueseq standard mRNA library kit (non-stranded, poly-A selection) following manufacturer’s guidelines. Libraries were then uniquely barcoded, pooled and sequenced on an Illumina Novaseq 6000 using single-end runs (at Weill Cornell Medical College’s Genomic Core Facility).

### RT- qPCR

Equivalent amounts of RNA were reverse transcribed using SuperScript III Reverse Transcriptase (Thermo Fisher Scientific) following manufacturer’s instructions. To normalize cDNA amount across samples, primers for *B2m* were used. cDNAs were mixed with gene specific primers and SYBR green PCR MasterMix (Sigma Aldrich) and run on an Applied Biosystems 7900HT Fast Real-Time PCR system.

### Single cell and bulk RNA-sequencing analysis

Trimmed FASTQ files were obtained from the Rockefeller University’s Genome Resource Center (scRNA-sequencing) or from the Genomic Core Facility (Weill Cornell Medical College; bulk RNA-sequencing), and raw sequencing reads were aligned to the mouse reference genome (UCSC release mm39) using STAR (v2.6)^79^. The expression values of each gene were quantified as both raw counts and transcripts per million (TPM) using Salmon (v.1.4.0)^80^, and compiled in R (v.3.6.1) using Tximport(v.1.12.3)^81^. *Bulk RNA-sequencing:* For differential gene expression analysis in R, low detection genes (minimum average read count <10) were filtered before DESeq2 analysis (v.1.16.1)^82^. Differential expression modelling used a negative binomial distribution and Wald test. Genes were differentially expressed for log_2_[fold-change]>|1| and adjusted *P*<0.05. Heatmaps and bar graphs illustrating differential gene expression were constructed in a Python environment (detailed in next paragraph).

#### scRNA-sequencing

Analysis and visualization of the data were conducted in a Python environment built on Pandas (v.2.0.1), NumPy (v.1.24.2)^83^, SciPy (v.1.10.1)^84^, scikit-learn (v.1.2.0), SCANPY (v1.9.3)^85^, AnnData (v0.9.1)^85^, matplotlib (v3.7.1)^86^ and seaborn^87^ packages. Raw count and metadata matrices for 1489 single outer root sheath cells across the hair cycle were loaded in SCANPY as an AnnData object. Single cell data was preprocessed to remove lowly detected genes (expressed in <75 cells) and cells with low complexity libraries (<2,000 genes detected). SCANPY was used to normalize counts per cell, and highly variable genes were detected. Prior to dimensionality reduction by principal component analysis (PCA), data was centered and scaled. PCA was performed on highly variable genes, with 100 components and the svd_solver using ‘arpack’ (SCANPY default setting). To construct a *k*-nearest neighbours graph on Euclidean distance, 41 principal components were used (which captured 25% of the variance in the data). Data was visualized using uniform manifiold projection (UMAP) in SCANPY, and clustering was done using the Leiden algorithm (with a resolution setting of 0.5). Cluster resolution was chosen after iterating through resolution parameters from 0.1 to 0.75, as best capturing both hair follicle cycle stages and anatomic location (upper bulge region/upper ORS versus hair germ/upper-middle ORS versus lower ORS). Marker gene expression based on the literature^10-12,75^, together with the FACS markers each population was sorted on, was used to identify clusters. SCANPY was used to visualize selected marker genes in dot plots, or as normalized counts visualized on UMAPs.

Differential gene expression based on cluster identity was used in DESeq2 to identify genes that varied as cells transitioned from late anagen growth phase to catagen. Differential expression was performed as described for bulk RNA-sequencing, with the modification of a threshold of 0.75 to construct Wald tests of significance. Gene set enrichment analysis (GSEA) on differentially expressed genes was performed using GSEA software (v.4.3.2)^88,89^, and run with the MSigDB 2022 mouse database. Gene set terms with FDR<0.1 and showing high normalized enrichment scores in catagen cells were considered interesting. To construct gene set scores based on the GSEA identified terms the corresponding *Mus musculus* gene lists were obtained by Amigo2 through the Gene Ontology consortium. The SCANPY “tl.score_genes” function was used to compute the average expression of each gene set across single cells, and normalized to a randomly sampled reference set of genes^90,91^. The resulting gene set scores were colour coded on corresponding UMAP visualizations of the data.

### ATAC-sequencing analysis

Trimmed FASTQ files were obtained from the Rockefeller University’s Genome Resource Center and aligned to the mouse reference genome (UCSC release mm39 using Burrows-Wheeler Aligner (BWA), using BWA-MEM with default parameters. The output .sam files were name-sorted and duplicate reads were marked and removed using SAMtools (v.1.17)^92^. Peaks were called on each replicate using MACS3 (v.3.0.0) using the callpeak command, BAMPE, and a mappable genome estimate of 1.87×10^9^ (from the ENCODE pipeline). The fraction of reads in peaks was calculated using bedTools (v. 2.31.0)^93^ and used to scale bigwig files equivalently in deepTools (v.2.0.0)^94^. Bigwig files were created from deduplicated, pooled replicate bam files using deepTools, and normalized as reads per genome coverage. Pooled replicate bigwig files were also used to calculate peak coverage matrices to plot heatmaps of centered differential peaks, extended by 1 kb upstream and downstream. Differential peak analysis was done in DESeq2, using read count matrices across each individual replicate from concatenated, merged union peak sets from each replicate. These union peak sets were created separately for *in vivo* samples and *in vitro* samples. Differential analysis used negative binomial modelling, and Wald’s test for significance. To assign peaks to nearest expressed gene, part of the Inferelator-prior (v.0.3.8)^95^ package was used. Peaks were assigned to genes if they fell 50 kb upstream or 5 kb downstream of the gene body and were curated for expression using either scRNA-seq (in vivo samples) or bulk RNA-seq (in vitro samples). To make sure that all potential enhancers for genes related to the apoptotic cell clearance program were identified, any unassigned intergenic peaks within approximately 200kb of phagocytosis-related genes were manually curated. If no genes were expressed transcriptionally in the interval between phagocytic gene and unassigned intergenic peak, the intergenic peak was considered a potential enhancer for said gene. Peaks of interest were visualized using the integrated genome viewer (IGV) software (v.2.13.2), together with .bed files of differential peaks.

Motif enrichment analysis for *in vivo* samples was performed in two ways: First, the MEME suite (v. 5.5.2) package XSTREME in web browser format was used to search for motifs enriched in differential peaks, using as background the union set of all peaks detected, and the JASPAR 2022 vertebrate CORE transcription factor motif database, with lengths of 6-18bp specified. Both known and *de novo* enriched motifs were collapsed to clusters based on similarity and ranked based on adjusted p-value. Second, the transcription factor occupancy prediction by investigation of ATAC-seq signal (TOBIAS, v.0.14.0)^96^ framework was used to perform chromatin footprinting analysis. Briefly, replicate-pooled bam files read coverage across the genome was calculated and corrected for Tn5 transposase cutting bias before footprint scores were calculated within the union set of called peaks. TOBIAS footprint scores were used to compute differential binding between anagen and catagen pooled replicates, or between *Rxra* WT and cKO pooled replicates. RXR-family catagen bound footprints were visualized in IGV by pooling each individual RXR-family member’s bed footprint file.

### Statistics

All data from every experiment were included for analysis unless an error was detected via failed positive or negative controls; in that case the entire experiment was excluded from analysis. Measurements were taken from distinct samples, unless stated otherwise. Experiments were not blinded, given the lack of ambiguity in phenotypes observed and internal controls used.

Statistical and graphical analyses were performed in Jupyter Notebooks, running a custom Python environment built as described in the single cell sequencing analysis section. Sample sizes, replicates and statistical tests used are indicated in each figure legend. Unless otherwise stated, unpaired two-tailed Student’s *t*-tests with a 95% confidence interval were performed to test for pair-wise differences among the means. Data is visualized as box-and-whiskers plots, with the box representing the first to third quartiles of the data set, the median line inside the box, and the whiskers extending a maximum of 1.5 times the inter- quartile range. Observations that fall outside this range are plotted independently. For clarity, each observation in a data set is also visualized as a point overlaid on the box plot.

## Acknowledgements

We thank E. Wong, M. Nikolov, J. Racelis, P. Nasseir, L. Hidalgo, M. Sribour, T. Omelchenk, L. Polak and G. Gray for technical assistance; S. Ellis, M. Laurin, S.Gur-Cohen, S. Lui, S. Baksh, R. Niec,, A. Gola., J Novak, P. Ghose, and O. Yarychkivska for discussions; S. Shaham for project advice; S. Mazel, S. Semova, S. Han and S. Shalaby at The Rockefeller University’s Flow Cytometry Core for conducting FACS sorting; C. Lai and S. Huang at the Genomics Core of The Rockefeller University; the Weill Cornell Medicine Genomics Resources Core Facility; The Rockefeller University Electron Microscopy Resource Center; and The Rockefeller University’s Bio-Imaging Resource Center. E.F. is a Howard Hughes Medical Investigator. K.S.S. is a New York Stem Cell Foundation-Druckenmiller Fellow and was the recipient of a Canadian Institute of Health Research postdoctoral fellowship, and a Rockefeller University Women & Science postdoctoral fellowship. K.A.G was supported by a Cancer Research Institute Carson Family Fellowship. S.Y. was the recipient of an F31 Ruth L. Kirschstein Predoctoral Individual National Research Service fellowship from the National Cancer Institute (NCI) and a Pilot Award from the Shapiro-Silverberg Fund at The Rockefeller University. M.T.T is supported by a National Institute of Arthritis and Musculoskeletal and Skin Diseases (NIAMS) 1K99AR079575-01A1 Pathway to Independence Award. A.R.B. was a Kenneth C. Frazier Fellow of the Damon Runyon Cancer Research Foundation (DRG: 2448-21). N.R.I. was the recipient of a NIAMS Diseases National Research Service Award (F31AR073110). C.J.C. is the recipient of a NCI F99/K00 pre to postdoctoral transition fellowship (F99CA264439). This study was supported by grants to E.F. from the National Institutes of Health (R01-AR050452 and R01-AR31737) and by The New York Stem Cell Foundation.

## Author contributions

K.S.S. and E.F. conceptualized the study, designed the experiments, interpreted the data, and wrote the paper. K.S.S. performed all experiments, with assistance from K.A.G, S.Y., M.T.T., A.B., Y.Y., N.I., and C.C. K.A.G, S.Y., M.T.T., and A.B. assisted with FACS experiments. K.A.G, S.Y., Y.Y., and N.I. prepared ATAC-seq libraries for sequencing. M.T.T. and A.B. collected skin samples for *Rxra* cKO-related experiments. K.S.S analyzed ATAC-seq data with input from Y.Y. K.A.G performed WT HFSC bulk RNA- seq. C.C. prepared RNA for bulk RNA-sequencing. J.M.L. performed all lentiviral injections. H.A.P. performed electron microscopy and interpreted images. C.V.R and S.G. contributed *Mertk* mice, and expertise. All authors provided input on the final manuscript.

## Data availability

All data supporting the findings of this study are available within the Article and its Supplementary Information. All single-cell and bulk sequencing data generated within this study have been deposited at the Gene Expression Omnibus (GEO) under accession code GSE230523.

## Code availability

Custom code for scRNA-seq and ATAC-seq for this study will be deposited at Zenodo. All other codes are available from the corresponding author on reasonable request.

## Supplementary Information

Supplementary information is available for this paper.

Correspondence and requests for materials should be addressed to Elaine Fuchs.

